# BioXTAS RAW 2: new developments for a free open-source program for small angle scattering data reduction and analysis

**DOI:** 10.1101/2023.09.25.559353

**Authors:** Jesse B. Hopkins

## Abstract

BioXTAS RAW is a free, open-source program for reduction, analysis and modelling of biological small angle scattering data. Here, the new developments in RAW version 2 are described. These include: improved data reduction using pyFAI; updated automated Guinier fitting and *D_max_* finding algorithms; automated series (e.g. SEC-SAXS) buffer and sample region finding algorithms; linear and integral baseline correction for series; deconvolution of series data using REGALS; creation of electron density reconstructions via DENSS; a comparison window showing residuals, ratios, and statistical comparisons between profiles; and generation of PDF reports with summary plots and tables for all analysis. In addition, there is now a RAW API, which can be used without the GUI, providing full access to all of the functionality found in the GUI. In addition to these new capabilities, RAW has undergone significant technical updates, such as adding Python 3 compatibility, and has entirely new documentation available both online and in the program.

## 1. Introduction

Small-angle solution scattering (SAS) of both X-rays (SAXS) and neutrons (SANS) is a popular structural technique for studying biological macromolecules. SAS provides information on the solution state of macromolecules and complexes, including, but not limited to, size and molecular weight, flexibility, degree of folding, and overall shape (Trewhella, 2022; Da Vela & Svergun, 2020; Brosey & Tainer, 2019; Meisburger *et al*., 2017; Jacques & Trewhella, 2010; Svergun & Koch, 2003). The growing popularity of SAS as part of the structural biology toolbox has many contributing factors: expanding data collection capabilities for more challenging systems, including the increasing number of hyphenated techniques like size exclusion or ion exchange chromatography coupled to SAXS (SEC-SAXS and IEC-SAXS) (Graewert & Svergun, 2022; Pérez *et al*., 2022); increasing automation of data collection and analysis to make the technique more accessible for new users (Tully *et al*., 2023; Rosenberg *et al*., 2022; Lazo *et al*., 2021); and an increasing awareness that SAS is highly complementary to other structural and biophysical techniques such as X-ray crystallography (MX), nuclear magnetic resonance (NMR), cryo-electron microscopy (cryoEM), and multi-angle (also called static) and dynamic light scattering (Trewhella, 2022; Brosey & Tainer, 2019; Grishaev, 2017). Additionally, SAS has proven an invaluable tool for studying intrinsically disordered proteins and liquid-liquid phase separating systems, which are not readily amenable to common high-resolution structural techniques such as MX and cryoEM (Lenton *et al*., 2022; Martin, Hopkins *et al*., 2021; Martin *et al*., 2020; Sagar *et al*., 2020; Riback *et al*., 2017; Kikhney & Svergun, 2015).

Providing a detailed review of SAS techniques and data analysis is beyond the scope of this paper (interested readers are directed to any number of excellent reviews, such as those cited previously). Briefly, data is collected as two-dimensional detector images, which are then radially averaged into one dimensional scattering profiles. Both the sample in solution and the solution by itself are measured, and the solution blank (‘buffer’) measurement is subtracted from the sample measurement to create a subtracted scattering profile (Skou *et al*., 2014). Sometimes, more advanced deconvolution methods must be used to determine the scattering profiles of overlapping components in the dataset (Tully *et al*., 2021). Model free analysis of the subtracted profile is then carried out – including a Guinier fit, calculation of molecular weight, creation of the pair-distance-distribution (P(*r*)) function by indirect Fourier transform (IFT), and creation of specific plots like the Kratky and Porod plots – to provide initial information about the sample. Based on the system, more advanced analysis can be carried out, including *ab initio* reconstruction of low resolution models, fitting high resolution models to the data, and modeling flexibility or polydispersity in the solution (Skou *et al*., 2014; Dyer *et al*., 2014).

There are a wide variety of software tools developed by the SAS community to carry out data reduction, analysis, and modelling, and we can provide only a non-exhaustive overview. The most popular software remains the ATSAS package (both desktop and web versions) (Manalastas-Cantos *et al*., 2021), which provides everything from initial reduction through advanced analysis and modelling, such as rigid body modelling with SASREF (Petoukhov & Svergun, 2005) and bead model reconstructions with DAMMIF (Franke & Svergun, 2009). ScAtter is another general-purpose analysis package (https://github.com/rambor/scatterIV).

Other data reduction options include pyFAI, DPDAK and Fit2D (Kieffer *et al*., 2019; Ashiotis *et al*., 2015; Benecke *et al*., 2014; Hammersley *et al*., 1996), along with beamline-specific processing pipelines (which may rely on one of the previous software packages to do the reduction) (Tully *et al*., 2023; Thureau *et al*., 2021; Cowieson *et al*., 2020) and software provided by laboratory x-ray source companies. Numerous tools are available for more specific types of analysis including (but not limited to): SASSIE, FoXS (and FoXSDock and MultiFoXS), WAXSiS, DADIMODO and BilboMD (all available as web servers) for testing and/or modelling high resolution data against SAS profiles (Curtis *et al*., 2012; Schneidman-Duhovny *et al*., 2016; Knight & Hub, 2015; Evrard *et al*., 2011; Pelikan *et al*., 2009); DENSS and Memprot for low resolution electron density or bead model reconstructions from SAS profiles (Grant, 2018; Pérez & Koutsioubas, 2015); and REGALS and US-SOMO for deconvolution of data (Meisburger *et al*., 2021; Brookes *et al*., 2016). Many other programs exist and readers are encouraged to do a literature search or consult with experts to find the best options for their particular analysis.

The BioXTAS RAW program is most similar to Primus (ATSAS package) (Konarev *et al*., 2003) and ScAtter, in that it provides a user-friendly GUI interface with both model independent and model dependent analysis, though unlike those programs RAW allows users to start with and reduce images to scattering profiles. Our focus in making RAW is to provide an open-source, free, easy to learn and easy to use program capable of doing the majority of what users need, including data reduction, producing subtracted profiles, carrying out standard analysis such as Guinier fits and generating P(*r*) functions, and some more advanced analysis such as shape reconstructions and deconvolution of overlapping chromatography coupled SAS data. RAW also provides a home for some advanced analysis techniques by directly incorporating (with permission, and often assistance, from the authors) techniques developed by others. This makes these techniques easier to use (e.g. by providing a GUI and documentation) and expands the number of users able to take advantage of these developments. Finally, RAW also incorporates some popular tools from the ATSAS package (separate installation required) for a more unified experience in RAW. Because one of our goals is for RAW to be an open-source toolbox for SAS, we have been working to provide open-source alternatives to all of the ATSAS tools we incorporate, and with the exception of theoretical profile calculation with CRYSOL this is now the case. Thus, even without ATSAS available, RAW provides a full range of reduction and analysis techniques for its users.

## 2. Program Overview

Many of the basics of the RAW program have been previously described (Hopkins *et al*., 2017; Nielsen *et al*., 2009), and this article focuses on updates to the program since the last publication. Here we provide a brief summary of the program and capabilities for readers not familiar with RAW. The main window of RAW is shown in Figure 1.

**Figure 1.**
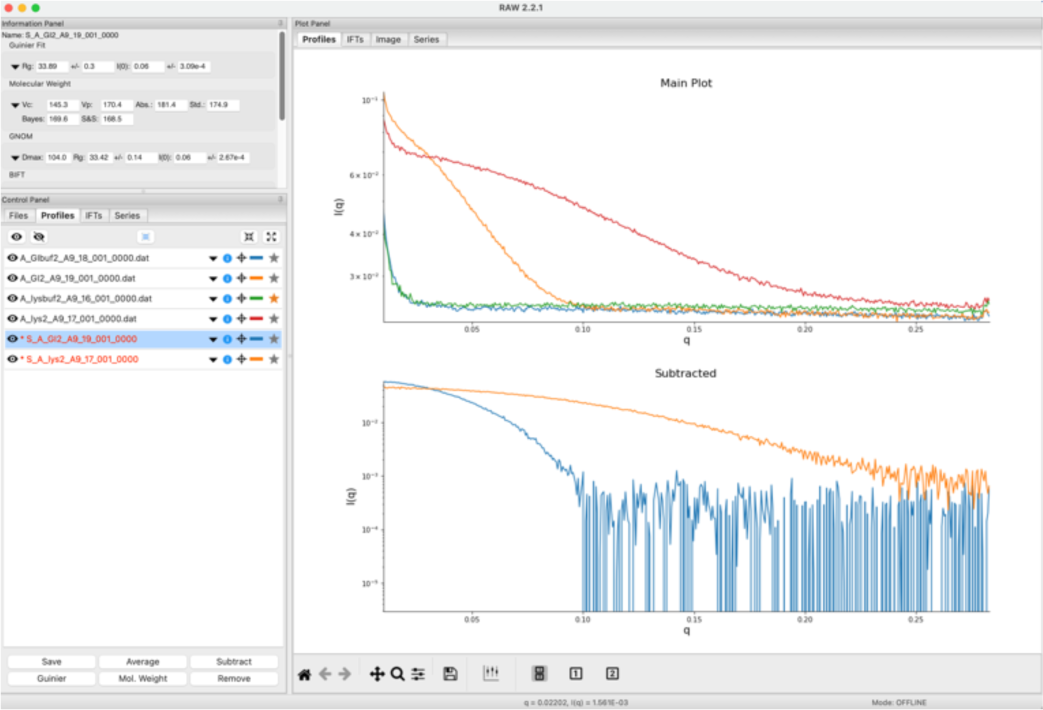
The main RAW window showing data for two batch mode protein datasets. The top plot shows the unsubtracted protein and buffer profiles while the bottom plot shows the subtracted protein profiles.

RAW is a GUI based, free and open-source Python program for reduction and analysis of small-angle solution scattering (SAS) data, both X-ray (SAXS) and neutron (SANS). The software is designed for biological SAS data. It is available on Windows, macOS (and OS X), and Linux. Using RAW, users can carry out analysis starting with detector images and go all the way through three dimensional reconstructions. They can mask, calibrate, and radially average x-ray images into one-dimensional scattering profiles as I(*q*) vs. *q*, where *q* = 4π sin(θ) /λ is the usual scattering vector with 2θ being the scattering angle and λ being the wavelength (note that RAW is ambivalent to the units of q, and users must label plots with the appropriate units, typically Å^-1^ or nm^-1^). RAW provides users with basic manipulation of scattering profiles including averaging, subtraction, rebinning, scaling intensity and *q* and trimming the *q*-range. Once a subtracted scattering profile is available, generated either in RAW or another program, there are a host of analysis and visualization tools available. Users can plot on linear and logarithmic axes in both intensity and *q*, generate Guinier, Kratky, and Porod plots, and create normalized and dimensionless Kratky plots (Durand *et al*., 2010). Users can perform Guinier fits to yield *Rg* and *I*(0). Molecular weight analysis is available to users by a number of methods: using a reference standard (Mylonas & Svergun, 2007), absolute scale calibration (Orthaber *et al*., 2000), the correlation volume (*Vc*) method (Rambo & Tainer, 2013), the adjusted Porod volume (*Vp*) method (Fischer *et al*., 2010; Piiadov *et al*., 2018), the ATSAS Shape&Size method (Franke *et al*., 2018), and the ATSAS Bayesian inference method (Hajizadeh *et al*., 2018). Users can generate P(*r*) functions using indirect Fourier transforms (IFTs) via either the ATSAS GNOM software (Svergun, 1992) or the Bayesian IFT (BIFT) method (Hansen, 2000).

After a P(*r*) function is available, either generated in RAW or in another program, RAW users can carry out ambiguity assessment of three dimensional (3D) reconstructions using the ATSAS AMBIMETER program (Petoukhov & Svergun, 2015). Users can then do 3D reconstructions, averaging, clustering, refinement, and alignment to high resolution structures within RAW using the ATSAS approach for bead models including DAMMIF, DAMMIN, DAMAVER, DAMCLUST, SUPCOMB/CIFSUP, and SASRES (Franke & Svergun, 2009; Svergun, 1999; Volkov & Svergun, 2003; Petoukhov *et al*., 2012; Kozin & Svergun, 2001; Tuukkanen *et al*., 2016). Alternatively, users can reconstruct electron density via the same steps minus clustering using DENSS (Grant, 2018).

RAW provides users with basic and advanced capabilities for dealing with liquid chromatography coupled SAS (LC-SAS) data and other sequentially sampled data. Users can load data as a series, and for standard size exclusion chromatography coupled SAS (SEC-SAS) a buffer region can be automatically selected and then subtracted from the dataset. *Rg* and MW are calculated across the elution peaks, and the peak region of interest can be automatically selected and the subtracted one-dimensional profile from that peak region generated for further analysis. RAW also provides users with the ability to carry out linear and integral baseline corrections to the data to account for various forms of background drift (Brookes *et al*., 2013, 2016). For more complicated series datasets, two deconvolution methods are available to users. Evolving factor analysis (EFA) deconvolution focuses mostly on SEC-SAS data or other data where components obey a first-in/first-out principle (Meisburger *et al*., 2016). Regularized alternating least squares (REGALS) provides a more general deconvolution approach that can apply additional constraints to the data such as smoothness or a real space restraint with a P(*r*) function (Meisburger *et al*., 2021). This makes REGALS more amenable to complicated datasets including titration series, time resolved measurements and ion-exchange chromatography coupled SAS (IEC-SAS).

In addition to the GUI version of the program, users can install RAW as a python package, giving access to the new RAW API, which they can use for custom python scripts, Jupyter notebooks, and in other programs (such as beamline processing pipelines) where they want to use the tools available in RAW in a more automated or scriptable fashion.

## 3. Availability

The RAW source code is GPL-3.0 licensed and available on GitHub: https://github.com/jbhopkins/bioxtasraw. Official versioned releases of RAW are made available for download on Sourceforge: https://sourceforge.net/projects/bioxtasraw/. Older releases of RAW can also be found on Sourceforge. These official versioned releases include the source code associated with that version as a .zip file, and prebuilt installers for Windows, MacOS (both x86_64 and arm64 native) and Linux (a .deb installer). Both the source code and installers are freely available for anyone to download, use, and share. RAW is also available through SBGrid. The RAW documentation is hosted on Read the Docs: https://bioxtas-raw.readthedocs.io/en/latest/.

## 4. Improvements to previously described features

### 4.1. Updates to Series data processing

The part of RAW that has undergone the most significant update since the last report on the program is the series processing capabilities. The core of the new features is a new LC Analysis module, Figure 2, that provides buffer subtraction, baseline correction, and sample range averaging all in one place. Additionally, there is a new deconvolution method for series data, REGALS, described in Section 5.3. One relatively minor but quite useful new feature is that users can now scale profile intensity and adjust the *q*-range of profiles for all the profiles in a series at once.

**Figure 2.**
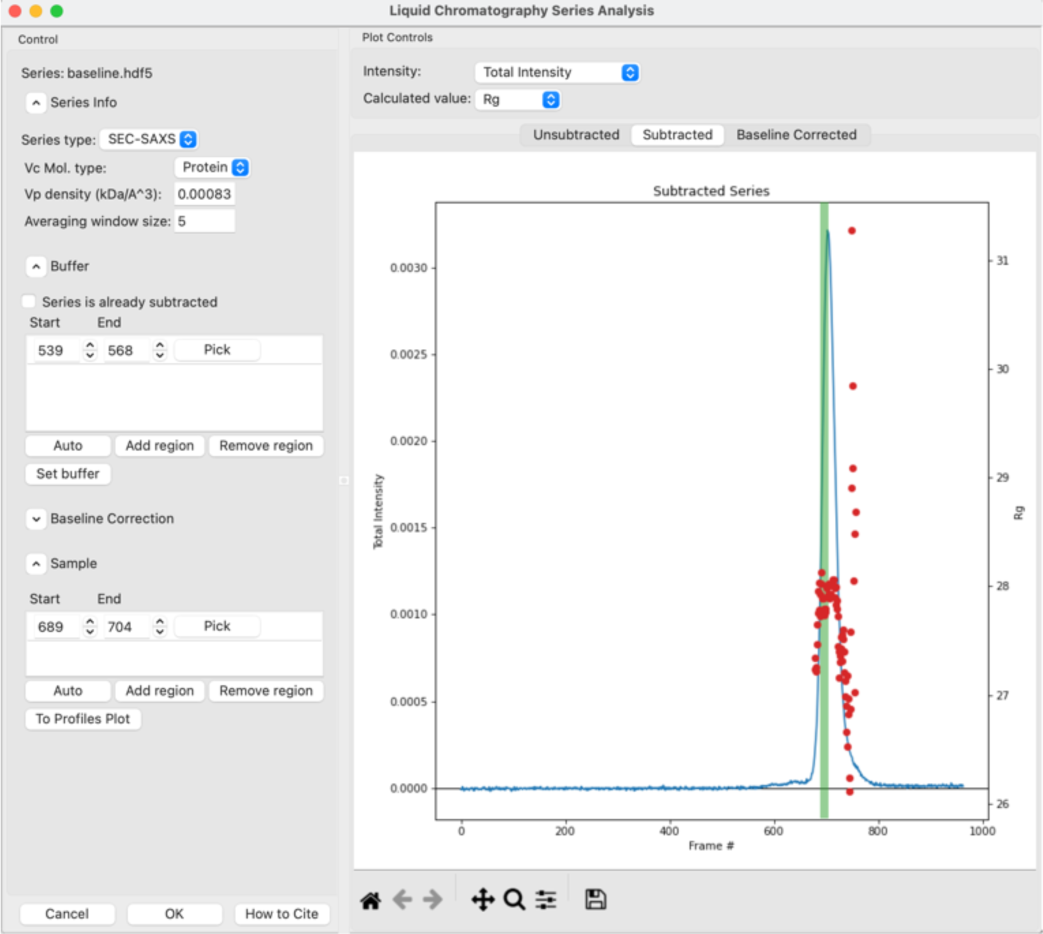
The new LC Series Analysis window in RAW, which allows automated and manual selection of buffer and sample ranges, baseline correction, and plots the *R_g_* and MW across the elution peak.

Before discussing the new features, it is useful to understand the general LC analysis workflow in RAW. First, a user will load in a series, which could be from .dat files, image files, or a previously saved RAW series file. The user then opens the LC analysis window. If any previous analysis has been done and saved, it is then displayed, if not the user proceeds as described below.

First, RAW displays a plot of the total intensity in each profile vs. frame number. The user then selects a buffer range (either automatically or manually, or a combination of both), which can be multiple disconnected regions of the data (e.g. pre- and post-peak buffer can be averaged). Once a user has selected a buffer range, RAW averages the buffer profiles and then subtracts this average buffer from every profile in the dataset. RAW then attempts to calculate *Rg* and MW for each profile in the dataset, using a sliding average window as previously described (Hopkins *et al*., 2017). The user sees a new plot of *Rg* or MW (user selectable) and the subtracted total intensity vs. frame number. The user may optionally then do a baseline correction (see Section 4.1.3). If they do, the correction is applied, *Rg* and MW are recalculated for the baseline corrected data, and the baseline corrected total intensity vs. frame number is plotted along with the new *Rg* and MW values on a new plot. The baseline itself is also plotted with the subtracted (but not baseline corrected) intensity for the user to examine.

The user next selects a sample range (either automatically, manually, or a combination of both), which can consist of multiple disconnected regions of the data, for further processing. Once the sample range is set, the original unsubtracted data is averaged across the selected sample range, and the averaged buffer profile is subtracted. This final subtracted profile is sent to the Profiles plot where the user may continue analysis. Note that this final subtraction step avoids averaging subtracted data where the same buffer profile has been used for subtraction of each profile, which would introduce correlations in the uncertainty values. The user can also work with previously subtracted data, skipping the buffer subtraction step. A more complete decision tree for standard SEC-SAS analysis is show in Figure 3.

**Figure 3.**
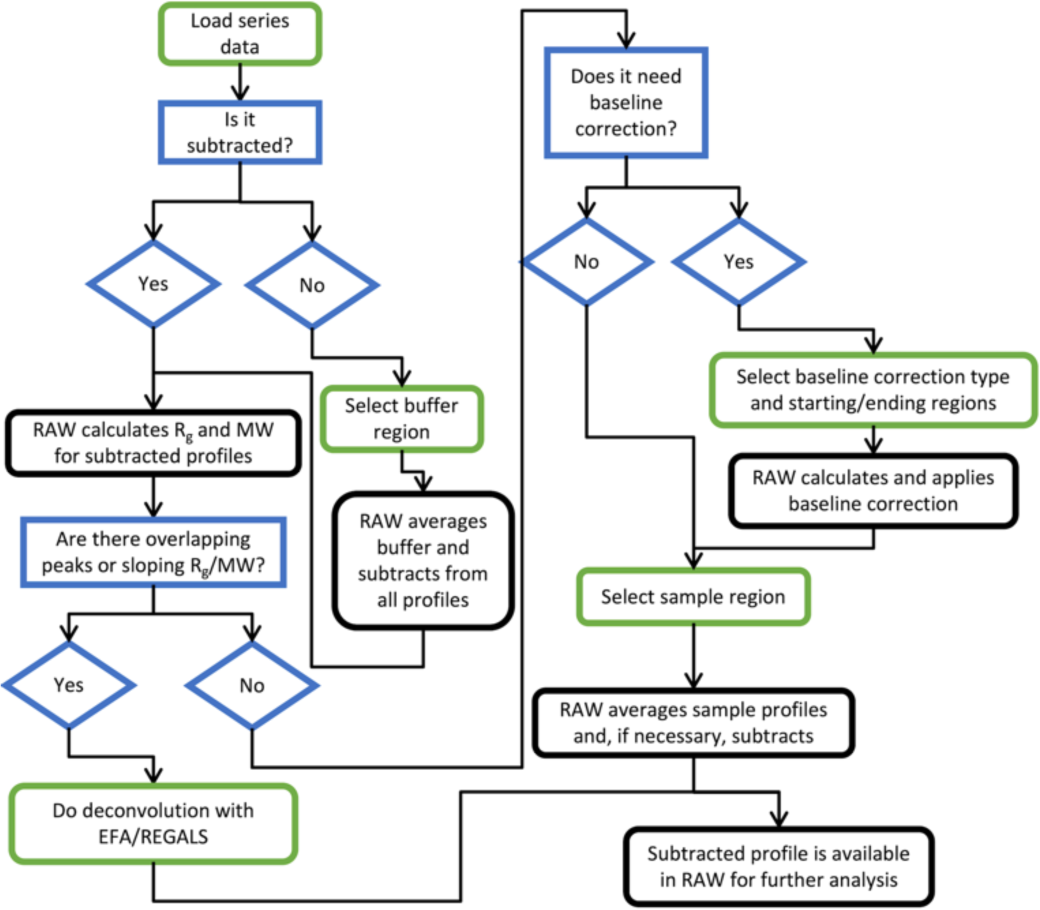
A decision tree for standard SEC-SAS analysis using RAW. Green edged boxes (rounded corners) indicate user actions (though many can be either manual or automated, such as picking a buffer region). Blue edged boxes (square corners) and diamonds indicate decisions, and black edged boxes (rounded corners) indicate processing done by RAW.

The selected buffer ranges, baseline correction parameters, and sample ranges are saved with the series data in RAW. If the LC Analysis window is reopened all the parameters are repopulated and can easily be tweaked by the user. These parameters are also saved with the profiles in the .hdf5 file when series data is saved, so data loaded into RAW from file has access to the previous analysis parameters.

In addition to the total integrated intensity, RAW can also display the mean intensity, the total intensity in a user defined q range, and the intensity at a specific q value. Different approaches to displaying the intensity have different use cases. The total intensity is the default choice in RAW because it provides a better view of possible issues with the dataset, such as being able to see capillary fouling or baseline drift at low or high q. Intensity in a specific q range (such as intensity from 0.02-0.2 Å^-1^, which is the default in CHROMIXS) can enhance the protein signal and make it easier to select peaks in low signal to noise datasets, but makes it easy for the user to overlook issues outside that q range. Generally speaking, for a well behaved and reasonable signal dataset, either total integrated intensity or intensity in a specific q range (that covers a reasonable range from low to mid q) yield similar plots. Users may change the displayed value to see what works best for their data set.

#### 4.1.1. New automated buffer and sample range selection

We have added the ability for RAW to automatically pick buffer and sample ranges from series datasets. This is aimed primarily at SEC-SAS datasets, but can be used for other series datasets with similar properties (i.e. distinct, constant buffer region and a distinct sample region). While this capability has been present in CHROMIXS (Panjkovich & Svergun, 2018) in the ATSAS suite for some time, we take a very different approach in RAW. For reference, the CHROMIXS approach automatically selects a sample range based on peak finding algorithms, often corresponding to the top of the strongest elution peak. Once the sample range is defined, a buffer range is selected by looking for contiguous low intensity ranges of the same length as the selected sample range. A subtracted scattering profile is created using the buffer range, and the buffer range is then scored based on the quality of the Guinier fit. The best quality range is selected as the buffer.

The automated selection in RAW proceeds in the opposite order. First a buffer range is selected, then a sample range is selected. The advantage to this order of operations is that additional information, such as trends in the *Rg* or MW across an elution peak, can be incorporated to ensure RAW picks an appropriate sample range in the dataset. The disadvantage to this order of operations is that RAW has less information for selecting the buffer range, and that long, low intensity flat regions that contain an elution component can accidentally be selected as the buffer range.

RAW’s basic approach to finding a good buffer range is to scan a window of defined size along the measured profiles, and test each range to see if it is a valid buffer range. If no valid range is found, the window size is narrowed and the scan repeated until either a valid range is found or the minimum size is reached. Additionally, RAW constrains the set of buffer ranges to test, both to avoid false positives and to improve the speed of the algorithm, based on the elution peaks in the dataset. A schematic of the search algorithm is given in Figure 4. A more detailed look at the algorithm and the tests used to determine if a buffer range is valid is provided in Section S1.

**Figure 4.**
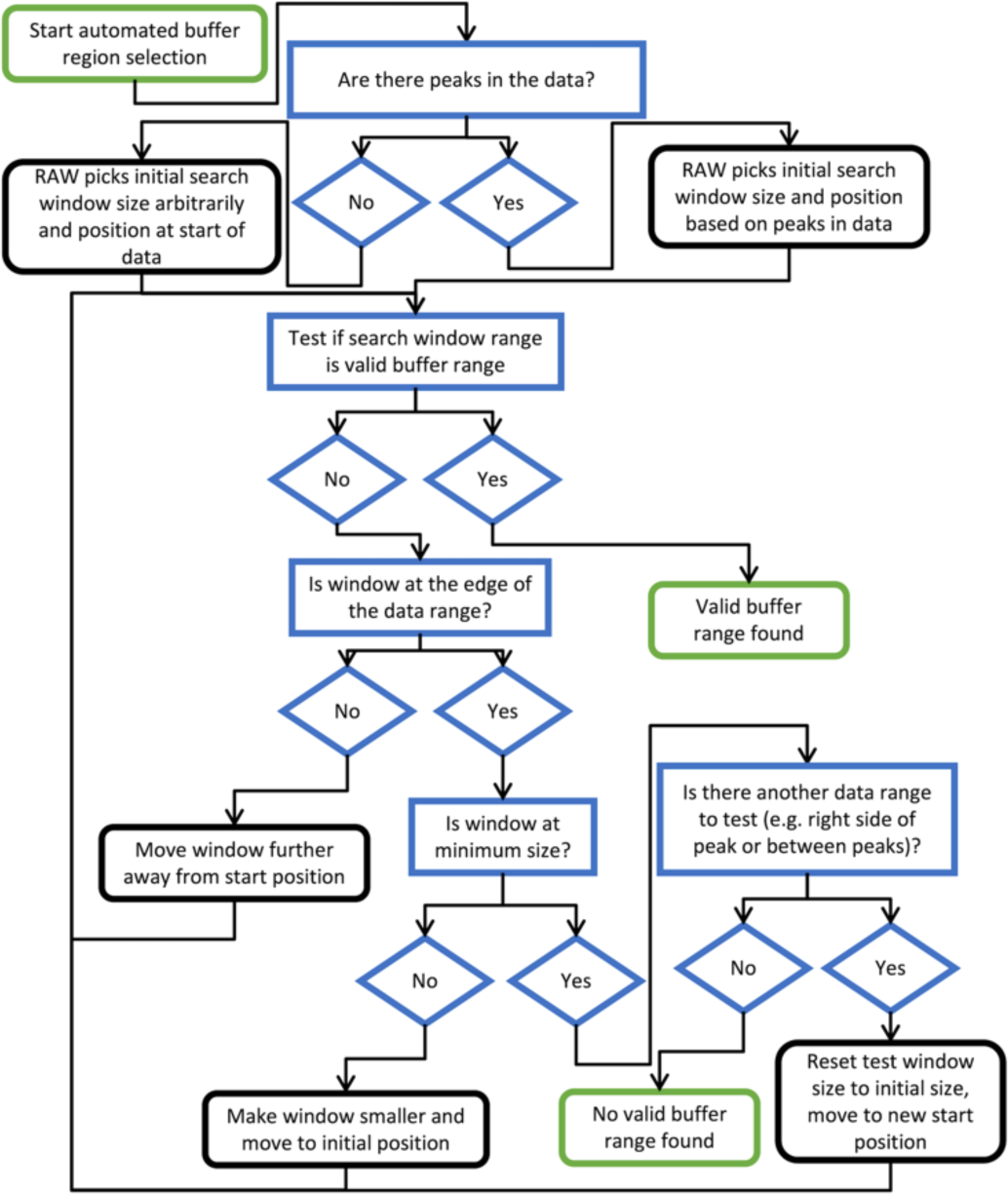
A flow chart for the automated buffer range finding algorithm used by RAW. Green edged boxes (rounded corners) are start and end points, blue edged boxes (square corners) and diamonds are decision points or tests in the algorithm, and black edged boxes (rounded corners) are actions by the algorithm.

The test for a valid buffer range has two inputs. The first is the total intensity (or mean intensity or intensity in a given *q*-range or at a particular *q* value, depending on user choice) vs. frames (or time) data, sometimes called the scattergram, and the second is the scattering profiles at each measured point in the elution. RAW evaluates a buffer range in three ways. First, it tests for correlations in the total intensity. Buffer scattering should have the same intensity at all measured points, so correlations are indicative of something eluting in the data (or an issue with the baseline). For the second test, RAW checks the similarity of the scattering profiles in the test range. We expect all buffer profiles to be similar. RAW checks similarity across three different *q* ranges: the low q range, where we may see capillary fouling or damage, the high *q* range, where we may see baseline drift, and the full profile. In the third test, RAW checks the number of significant singular values in the scattering profiles in the range. Buffer ranges should only have one significant singular value, the buffer scattering component. If any of these tests fail, the range is not a valid buffer range.

In order to optimize the speed of the automated buffer finding, the RAW runs the parts of the test in the order listed above, from fastest to slowest, and if any part fails on the selected range RAW does not run the subsequent parts.

RAW uses the same general approach for the automated sample range determination as it does for the automated buffer range finding, a window is scanned along the data and it tests whether each selected range is a valid sample range. If no valid range is found, the window size is narrowed and the scan repeated until either a valid range is found or the minimum size is reached. RAW again constrains the sample ranges to test to be within the strongest elution peak, both to avoid false positives and to improve the speed of the algorithm. A valid sample range will not be found if no peaks are found in the dataset. A schematic of the search algorithm is given in Figure S1. A more detailed look at the algorithm and the tests used to determine if a buffer range is valid is provided in Section S1.

The test for a valid sample range has three inputs: the scattering profiles and the *Rg* and MW calculated for each profile in the selected range. There are five parts to the test. The first part is simply whether all profiles in the selected range have calculated *Rg* and MW values. If the selected range is good, these values should be calculated for all profiles in the range. The second part checks for correlations in the *Rg* and MW values in the range. If the sample is uniform across the selected range there should be no correlation. The third part tests for similarity between the subtracted scattering profiles in the selected range. We expect all subtracted profiles to be similar to within a scale factor. As with the buffer similarity test, the full *q* range, the low-*q* range, and the high-*q* range are all tested. For the fourth part, RAW checks the number of significant singular values in the scattering profiles in the range. As with the buffer range, sample ranges should only have one significant singular value, the sample scattering component. The fifth and final part of the test is to check whether including all the profiles in the selected range improves the signal to noise of the final averaged subtracted scattering profile. If including a profile in the average decreases the signal to noise, then that profile should not be included in the final dataset and so the selected range is not valid. If any of these tests fail, the range is not a valid sample range.

As with the automated buffer range selection, in order to optimize the speed of the algorithm RAW runs the tests in the order listed above, fastest to slowest, and if any test fails on the range subsequent tests are not run.

It should be noted that both the buffer and sample range finding algorithms in RAW are simply based on useful heuristics. There is no proven way of always finding valid sample and buffer ranges, this may not exist, so instead we have devised a set of metrics that corresponds to how a well-trained human would evaluate the data (e.g. for the sample, it should have constant *Rg* and MW, the profiles should be the same, and we shouldn’t include data that’s too noisy) and which yields results that consistently pass the eye test from users. The algorithms can yield invalid ranges, and so the user should always examine and evaluate the automatically selected ranges before proceeding with further analysis, and modify them as necessary.

We show a comparison of automatically determined buffer and sample ranges using RAW and CHROMIXS (from ATSAS 3.2.1) in Figure 5 (data for this figure are from proteins measured by users at the BioCAT beamline, used with permission, and the data is anonymized. A general data collection protocol is given in Supporting Information S3. For high quality data, Figure 5a, both programs yield similar, overlapping, sample ranges. Because the *Rg* calculated across the peak tails off slightly at the edges (starting near ∼21.3 Å on the left edge, plateauing near ∼21.6 Å in the middle, and dropping to ∼20.8 Å on the right edge) RAW picks a more conservative range of the peak than CHROMIXS does, avoiding this tailing. There is a significant difference in selected buffer range, but as the buffer is uniform this results in no appreciable difference in the subtracted profile. In this case, the final profiles (not shown) are essentially identical, though the CHROMIXS one has slightly better signal to noise due to selecting more of the peak.

**Figure 5.**
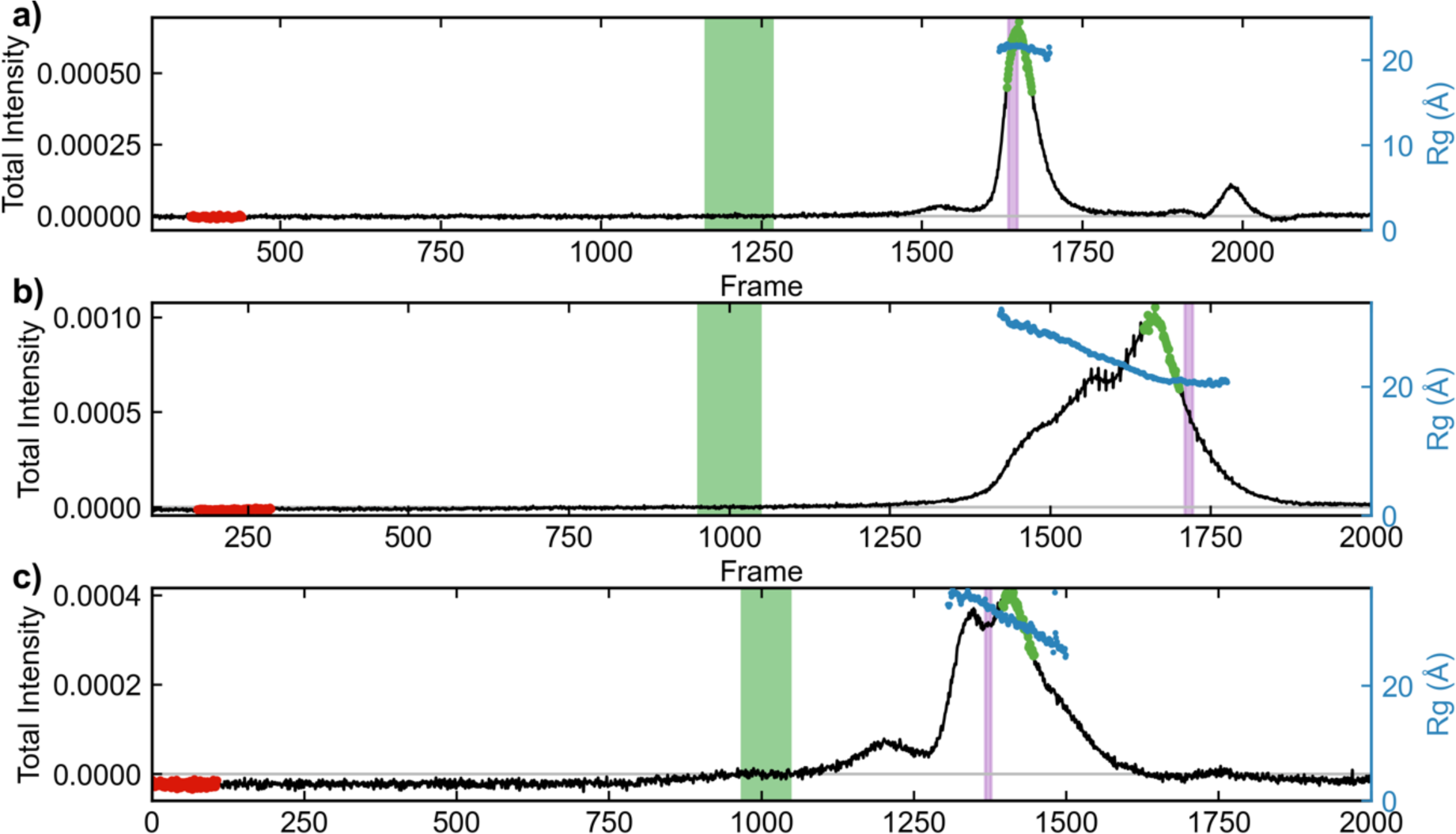
Automated buffer and sample regions as selected by RAW (green and purple shaded regions) and CHROMIXS (red and green dots) for various SEC-SAXS datasets. The total subtracted intensity for each dataset is shown in black (left axis), and the calculated *R_g_* across the elution in blue (right axis). Subtracted profiles were calculated using the buffer range selected by RAW, and *R_g_* calculations were done using RAW. The plots show: a) Well behaved SEC-SAXS data, showing a flat buffer region, a well separated single peak, and relatively constant *R_g_* across the peak. The sample ranges selected by RAW and CHROMIXS are overlapping, with CHROMIXS selecting a larger region while RAW excludes frames where the *R_g_* starts to trend slightly downward on the edges of the peak. b) A poorly separated sample showing only a small region of potentially usable constant *R_g_* profiles on the right edge of the peak. Here RAW selects frames in that constant *R_g_* region, where CHROMIXS fails to do so and selects frames in the obviously overlapped region where the *R_g_* trends upwards. c) A rare case where RAW selects a buffer range obviously within the elution region, resulting in incorrect background subtraction. CHROMIXS selects an appropriate buffer range.

Several other datasets were selected to highlight where the algorithms can break down. In Figure 5b, the elution profile and sloping *Rg* clearly show there are multiple overlapping components in the elution. Because it has access to the *Rg* information, RAW selects a sample range near the trailing edge of the peak where the *Rg* is essentially constant. CHROMIXS selects a sample range covering a large portion of the peak where the *Rg* varies. The resulting profiles (not shown) are not the same, and the profile from RAW is of higher quality. While in cases like this we would recommend that users apply a deconvolution technique such as EFA rather than averaging ranges of the peak, the additional information available to RAW clearly generates a better result. We haven’t tested this kind of analysis extensively with CHROMIXS, so we cannot say how common an occurrence this inaccurate sample range selection is when there are multiple components in solution.

Similarly, RAW is not immune to mistakes. In Figure 5c, RAW selects a buffer range that meets all the automated criteria (flat range, similar profiles, earlier than the first identified peak), but which is clearly in a region where protein is eluting. CHROMIXS, presumably because it includes the additional constraint of the best Guinier fit, avoids this and selects an earlier buffer range (note that in this elution there is no good range to pick for the sample, as seen by the continuously sloping *Rg*, but both algorithms return a result. A user should instead apply a deconvolution approach). Again, we see a trade-off based on where one includes additional information. In our anecdotal (though extensive) experience, failures of this type are rare for RAW, and we believe the trade-off of this failure mode vs. being able to determine what portions of the peak contain multiple components based on changes in the *Rg* and MW is a valuable one.

#### 4.1.2. Testing and reporting on suitability of selected buffer and sample ranges

Whenever the user sets the buffer or sample range in the LC Analysis panel, RAW validates the range using the tests described above. If any of the tests return an invalid result, a summary dialog of the failed test results is shown to the user and the user is given the choice to continue with their selected range or to adjust the range. While it is often acceptable to include ranges with reported validation issues (particularly for the sample range there is often only a small range that is completely valid, so accepting some potential issues to improve the overall data quality can be a reasonable choice), the user is at least made aware of the issues and can decide for themselves if they want to proceed.

#### 4.1.3. Series baseline correction

We have added two baseline correction algorithms to RAW to correct for drift or damage, a linear baseline correction that is useful for simple drift (such as from changes in temperature), first described in (Brookes *et al*., 2013), and an integral baseline correction that is more useful to account for damaged species (sample or buffer) accumulating on the capillary (Brookes *et al*., 2016). In both cases the user first subtracts the data, then defines start and end ranges for the correction. RAW averages the profiles in the start and end ranges to create the average start and end points used for the correction. The user can select whether the correction should be applied to just the data between the start and end points or all the data in the series. The linear correction is applied independently to each *q* value in a profile. In brief, for every *q* value a simple linear fit in intensity between the start and end points is carried out, and then the intensity of that fit line at a particular profile is calculated, and subtracted from that profile. The integral baseline correction uses the algorithm previously described and implemented in US-SOMO (Brookes *et al*., 2016).

#### 4.1.4. New .hdf5 save format for series

Previously RAW saved series data as a “.sec” file. While technically an open format, in that anyone could look up the specifications for reading or writing in the RAW code, it relied on the Pickle module in python to serialize and compress the data, which meant that only other python programs could open these files. We have changed RAW to use HDF5 files to save series data, which is a more standard format that can be opened by any program with HDF5 compatibility.

### 4.2. Improvements to automated Guinier fitting and *Dmax* finding

The new version of RAW provides an updated heuristic algorithm for determining the range of a Guinier fit (“auto Guinier”) and a new heuristic algorithm for finding the optimal maximum dimension (*Dmax*) for a P(*r*) function (“auto *Dmax*”) being calculated with an indirect Fourier transform (IFT). In both cases, the algorithms were developed and tested against the data available in the SASBDB in fall 2020 (Kikhney *et al*., 2020).

To demonstrate the utility of these automated methods, and to compare against other available methods, we compared the results against values determined by scientists from real experimental data. To this end, we ran the methods against all available biological macromolecular data in the SASBDB (as of June 2023). We used a simple python script to scrape the SASBDB using the provided REST API and download the scattering profiles and the depositor reported *Rg* and *Dmax* values for all biological macromolecular entries. We only used entries with a molecule type listed in the SASBDB of Protein, RNA or DNA, a total of 3138 datasets, treating the *Rg* and *Dmax* values reported by the depositor/experimenter as the “true” values for the dataset. We used the RAW auto Guinier and *Dmax* algorithms from version 2.2.0 of RAW; ATSAS programs were from ATSAS version 3.2.1.

#### 4.2.1. Automated Guinier fitting

The basic implementation of the auto Guinier function was described previously (Hopkins *et al*., 2017). We made two major changes in this updated version. First, some of the weighting coefficients on the different component tests were changed to yield better results based on tests against the SASBDB data. Second, the algorithm now undergoes a progressive relaxing of certain criteria and changing of the weights, which allows automatic determination of the Guinier range from lower quality (noisy, aggregated, etc) data. Here, RAW first does a relatively strict search, looking for only high-quality regions. If that fails to yield a suitable region, RAW allows a smaller window for the fit and lower the minimum acceptable quality of a fit. If that fails, RAW then allows the fit to include more data, both by broadening the region of the profile over which the search takes place and by increasing the allowable range for minimum and maximum *qRg* values, RAW also further reduces the quality threshold for a successful fit. These changes provide a much more robust function for low quality data and yield high quality results, as discussed below.

We ran the RAW auto Guinier algorithm and the ATSAS AUTORG program (Petoukhov *et al*., 2007) on the collected SASBDB profiles to obtain Guinier fits. If the experimenter reported starting and ending *q* values agreed precisely with either of the automated results we did not include the dataset in subsequent evaluation, as the experimenter may have simply reported the value from an automated method and we wanted only to compare to manually determined values. We used the ratio of the automated values and the true *Rg* to determine how close each automated determination was to the “true” value. We also tracked the number of datasets where a method failed to return a value. Of the 3138 initial datasets, 1827 had values that didn’t match either of the automated methods tested and had experimenter provided *Rg* values. From these, the average and standard deviation of the ratio of (experimenter determined *Rg*)/(automatically determined *Rg*) was 1.04 ± 0.56 for the RAW auto Guinier method and 1.03 ± 0.63 for the ATSAS AUTORG method. Plots of the automated *Rg* vs. experimenter determined *Rg* are shown in Figure 6. RAW failed to return results for 5 (0.27%) datasets and AUTORG failed to return results for 11 (0.60%) datasets. Our results show that both algorithms are robust, in that they fail on less than 1% of all datasets, and that both algorithms are on average quite accurate. For comparison, the old RAW algorithm, run on the same set of data, was similarly accurate (*Rg* ratio: 1.02 ± 0.51), but failed on a significant number of datasets (181, 9.9%). When we used all datasets, including those where the experimenter input *q* range matches that determined by one of the automatic methods, there was no significant change in the results (see Supporting Information S2).

**Figure 6.**
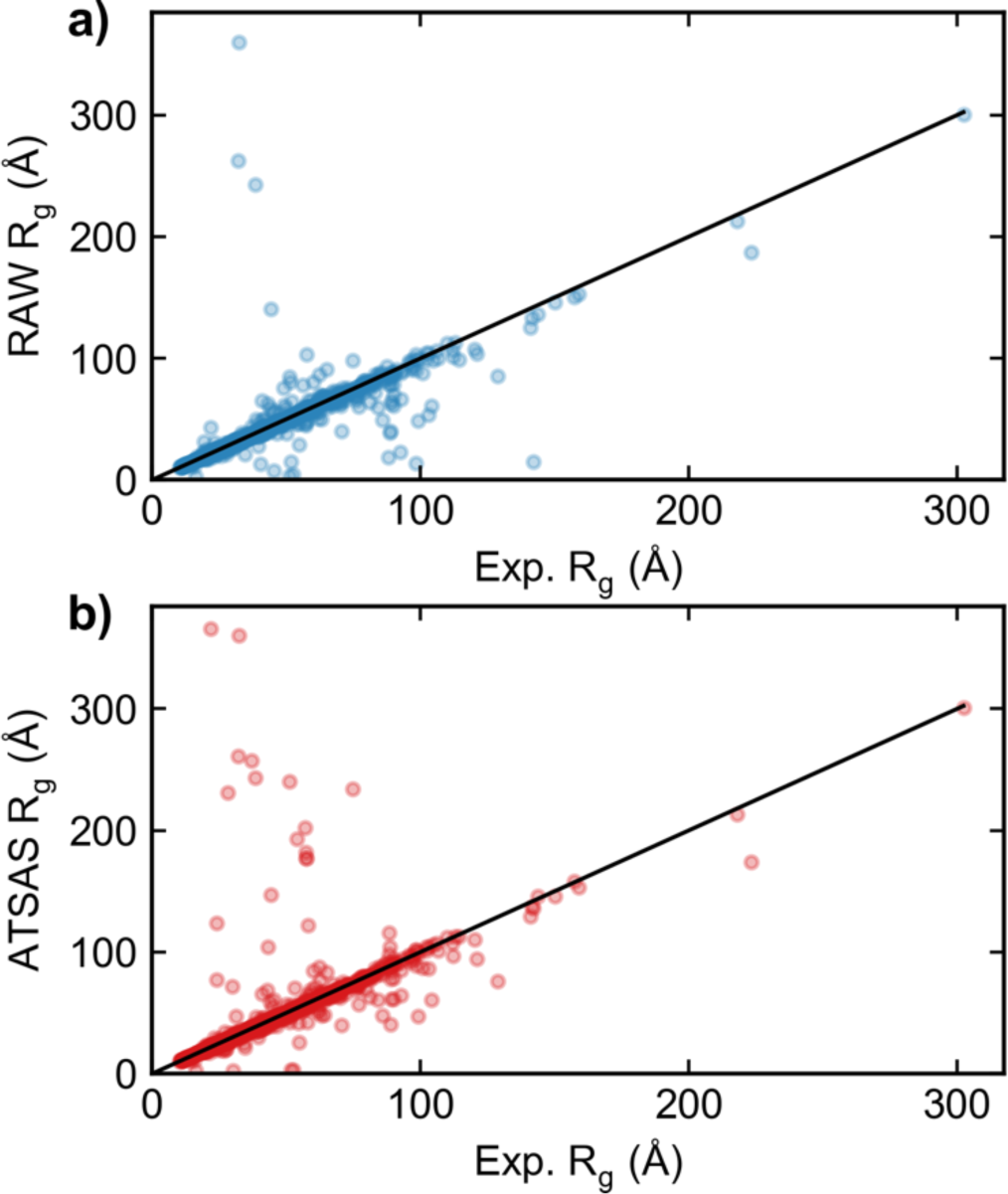
Plots of the automatically calculated *R_g_* values by a) the RAW automatic Guinier function, and b) the ATSAS AUTORG function on the y-axis vs. the experimenter determined *R_g_* from a SASBDB entry on the x axis. Results are shown for all SASBDB entries with *R_g_* values that were classified as either Protein, DNA, or RNA, unless the experimenter determined Guinier range perfectly matched either automated method. Perfect agreement between the automated method and the experimental method would be equal *R_g_* values, shown by the black line in each figure.

### 4.2.2. Automated *Dmax* determination

The auto *Dmax* function can run in several ways. If the ATSAS package is not available, it simply returns the *Dmax* value found by BIFT. However, if the ATSAS package is available then *Dmax* can be fine-tuned to get a more accurate value. The basic idea is simple. First, RAW runs other automated methods – BIFT, DATGNOM (Petoukhov *et al*., 2007) and DATCLASS – to determine a good starting point for the search. After determining a starting value, RAW calculates the P(*r*) function using GNOM with force to zero at the maximum dimension turned off. RAW checks this initial unconstrained P(*r*) function in two ways, first for overestimated *Dmax* values then for underestimated *Dmax* values, and adjusts the maximum value until it finds a good *Dmax*.

In a P(*r*) function with an overestimated *Dmax*, we expect either a long tail oscillating about zero (for homogenous, monodisperse, non-interacting samples) or negative values near the maximum dimension (for data with repulsive interactions) (Jacques & Trewhella, 2010). Using these criteria, if RAW determines that *Dmax* is overestimated, it decreases *Dmax* in 1 Å increments and recalculates the unconstrained P(*r*) function using GNOM until these criteria are no longer satisfied. In a P(*r*) function with an underestimated *Dmax*, we expect that the value at the end of the P(*r*) function is significantly greater than zero (Jacques & Trewhella, 2010). Using these criteria, if RAW determines that *Dmax* is underestimated, it increases *Dmax* by 1 Å and recalculates the P(*r*) function until that is no longer the case. More details on how these criteria are applied to determine under and overestimation are in Section S2.1. RAW applies one additional constraint to the adjustments, constraining the change in *Dmax* to be no more than 50%, either an increase or decrease, of the initial value, to prevent the algorithm from running away. This is particularly useful in the cases of highly aggregated data where there may be no appropriate maximum dimension and the algorithm could otherwise increase *Dmax* essentially indefinitely.

When taken together, these two simple adjustments for overestimated and underestimated *Dmax* values provide a more robust estimate of the maximum dimension than any of the other tools mentioned above, though we rely on those tools to find an appropriate starting point and so our approach should be considered complementary to the previously developed methods.

To test the accuracy of the automated *Dmax* algorithms, we ran the RAW auto *Dmax* algorithm, the BIFT function, and DATGNOM and DATCLASS programs in the ATSAS package on the collected SASBDB profiles to obtain *Dmax* values. As with the *Rg* tests, if the reported *Dmax* values agreed precisely with any of the automated values (rounded to the integer precision reported in the SASBDB), we did not include the dataset in subsequent evaluation. We calculated the ratio of the experimenter reported *Dmax* and the automated values to determine how close the automated methods were to the “true” value. We also tracked the number of datasets where a method failed to return a value. Of the 3138 initial datasets, 2502 of them had values that did not match any automated method and had experimenter provided *Dmax* values. Table 1 shows the average and standard deviation of the ratio of (experimenter determined *Dmax*)/(automatically determined *Dmax*) for the various methods, as well as the number of datasets where the algorithm failed to return a result. Plots of the automated *Dmax* vs. experimenter determined *Dmax* are shown in Figure 7. If we use all datasets, including those where the user input *Dmax* matches that determined by one of the automatic methods, there was no significant change in the results (see Supporting Information S2.1). There are several interesting takeaways here. First, none of the automated *Dmax* methods are as good as either of the automatic Guinier determination methods, showing that automatically determining *Dmax* is still a challenge. This may well reflect the inherent uncertainties in the IFT method. Second, most of the methods tend to have a systemic bias, either towards underestimating or overestimating the *Dmax*. In particular, RAW and BIFT both overestimate, RAW by about 11% and BIFT by about 15%, while DATGNOM tends to underestimate by about 25%. Of the non-RAW methods, DATCLASS is the closest on average to the experimental value. However, DATCLASS also fails for 20% of the datasets, while none of the other methods had any failures. Overall, the RAW auto *Dmax* is the best combination of robustness and accuracy, although this result is likely expected since we take the other three methods as an input and then refine.

**Table 1.** The average and standard deviation of the ratio of (experimenter *D_max_*)/(auto *D_max_*) for different automated *D_max_* determination methods used on the biological data in the SASBDB. The number of failures for each method is also reported.

| Algorithm | Mean ratio ± std. | Number of failures |
| --- | --- | --- |
| RAW auto $D_{max}$ | 0.89±0.51 | 0 (0%) |
| BIFT | 0.84±0.69 | 0 (0%) |
| DATGNOM | 1.27±0.98 | 0 (0%) |
| DATCLASS | 1.06±0.56 | 501 (20.0%) |

**Figure 7.**
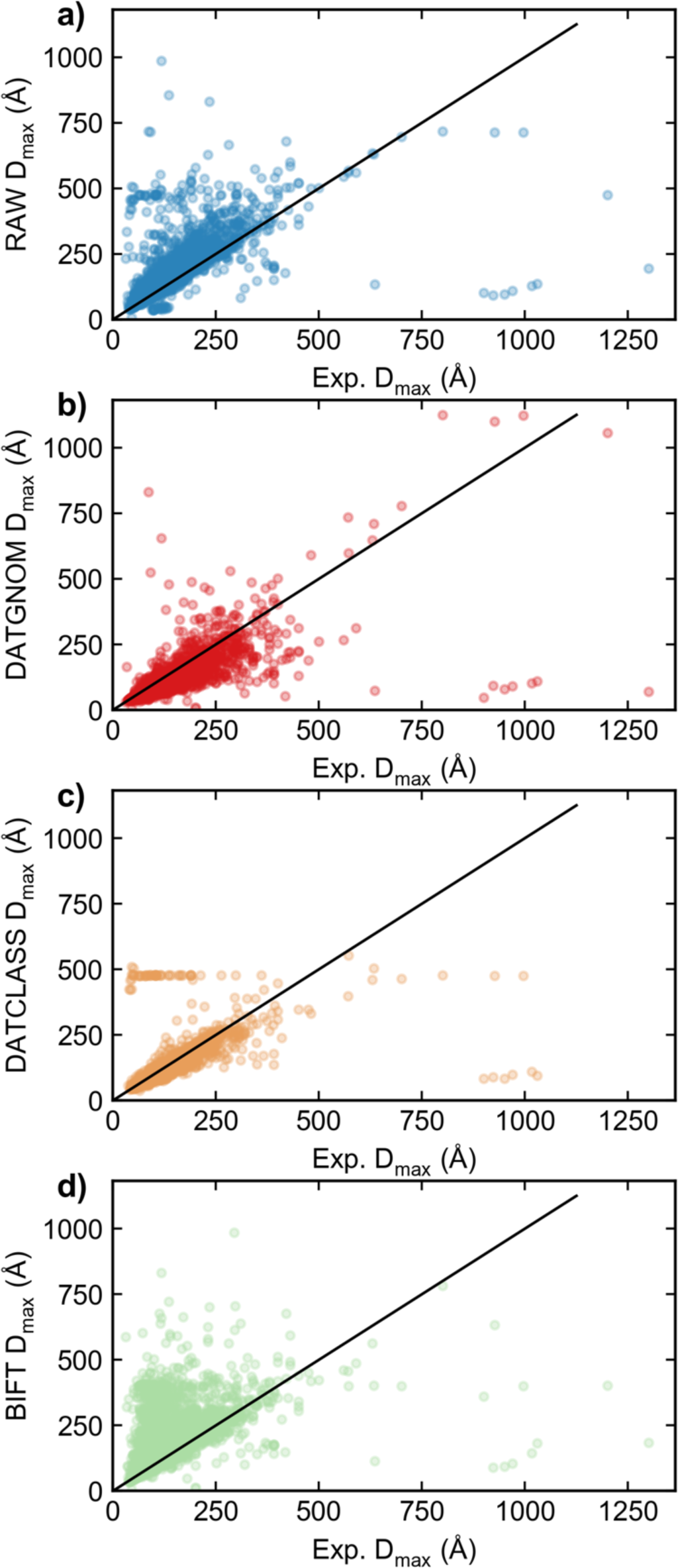
Plots of the automatically calculated *D_max_* values by a) the RAW auto D_max_ function, b) the ATSAS DATGNOM function, c) the ATSAS DATCLASS function, and d) BIFT (as implemented in RAW) on the y-axis vs. the experimenter determined *D_max_* from a SASBDB entry on the x axis. Results are shown for all SASBDB entries with *D_max_* values that were classified as either Protein, DNA, or RNA, unless the experimenter determined *D_max_* perfectly matched any automated method. Perfect agreement between the automated method and the experimental method would be equal *D_max_* values, shown by the black line in each figure.

### 4.3. Improved uncertainty estimation for Guinier fits

Uncertainty estimation in values derived from the Guinier fit, particular the *Rg* and *I*(0) values, is challenging because there are two competing sources of uncertainty. First, there is the inherent uncertainty in carrying out the fit, which is easily captured by the covariance of the fit parameters (assuming accurate uncertainty estimates for the intensity values of the scattering profile). Second, there is the uncertainty in selecting the range of data to be fit, which is not easy to quantify. We are certainly not the first to acknowledge this challenge, for example in the ATSAS package the uncertainty in the *Rg* and *I*(0) values from the AUTORG (Petoukhov *et al*., 2007) function, which automatically determines the endpoints is different (and typically larger) than the uncertainty provided using the DATRG function with the endpoints determined by AUTORG.

In order to provide more robust uncertainty values from the Guinier fit, we created an algorithm that estimates the effect of changing the endpoints of the Guinier range. RAW assumes that the endpoints should not be extended, that the user has picked the start and endpoints such that the data outside the selected range is not usable (for example, the start point may be picked to exclude aggregation or repulsive effects, while the end point is typically picked at the largest *q* value where the Guinier approximation still holds). RAW generates a series of sub ranges contained within the user selected Guinier range. It then calculates the *Rg* and *I*(0) values for Guinier fits to these sub ranges. RAW takes the standard deviation of these values and provides that as an additional uncertainty estimate, the range induced uncertainty. RAW shows a top-level uncertainty value in the Guinier fit window that is the greater of either the fit uncertainty or the range uncertainty, and the individual values are also provided so the user can decide which they wish to use.

For well-behaved data and well selected Guinier ranges, the largest source of uncertainty is typically from the covariance of the fit parameters. However, for data which displays systematic non-linearities (due to the presence of aggregation, repulsion, or a poorly selected fit range that extends past a *q*-range where the Guinier approximation is valid) the range uncertainty is often the largest uncertainty. For example, using data from the RAW tutorial, the provided glucose isomerase data is high quality, and returns an *Rg* of 33.89 Å over the range determined by RAW’s automated Guinier fit function. For this data, the fit uncertainty is 0.24 Å, and the range uncertainty is 0.10 Å. In this case, the data all falls on the expected Guinier fit line, and so the uncertainty from the measured intensity, which results in the fit uncertainty, is larger than that of the choice of fit endpoint. The tutorial data also includes highly repulsive data from concentrated lysozyme. The reported *Rg* here is 12.18 Å, with a fit uncertainty of 0.015 Å and a range uncertainty of 0.18 Å, indicating that because there is a repulsive effect the choice of range introduces much more variation in the Guinier fit values than the uncertainty in the measured intensity.

If the Guinier range is determined using RAW’s automated method, instead of showing the range uncertainty described above, RAW provides a similar value, the standard deviation of the *Rg* and *I*(0) values of all ranges found during the automated search that exceed a specified quality threshold (analogous to the uncertainty provided by the ATSAS AUTORG function).

### 4.4. Handling of multi-image files

Many modern detectors, such as the commonly used EIGER (Dectris) detectors, use file formats that package multiple images into a single file (e.g. an HDF5 file). We have updated RAW to be able to efficiently load in and radially average multi-image files, including the ability to determine the appropriate file numbering for series data loaded from several multi-image files, as well as display images from these files. The FabIO python library (Knudsen *et al*., 2013) provides the basic reading of the file formats, while RAW deals with which parts of the file to process or display.

### 4.5. Use of pyFAI for radial averaging

Since the last report about RAW, we have modified RAW to use the pyFAI python library (Ashiotis *et al*., 2015; Kieffer *et al*., 2019) for radial averaging as well as detector calibration. The major advantage of this change is that pyFAI is extremely fast and can utilize GPUs, so that even very large detector images can be processed quickly. pyFAI also incorporates additional corrections not available in the previous RAW radial averaging algorithm, such as correcting for incident beam polarization direction. This change has allowed RAW’s radial averaging to keep up with the demands of modern data collection, both in terms of image size and frame rate.

### 4.6. ATSAS Integration

RAW provides an interface for several programs from the ATSAS package (separate installation required) (Manalastas-Cantos *et al*., 2021), including: AMBIMETER (Petoukhov & Svergun, 2015), CIFSUP (ATSAS version 3.1.0 or newer), CRYSOL (Svergun *et al*., 1995; Franke *et al*., 2017), the Bayesian inference MW method in DATMW (Hajizadeh *et al*., 2018), DAMMIF (Franke & Svergun, 2009), DAMMIN (Svergun, 1999), DAMAVER (Volkov & Svergun, 2003), DAMCLUST (prior to ATSAS version 3.1.0) (Petoukhov *et al*., 2012), DATCLASS (Franke *et al*., 2018), GNOM (Svergun, 1992), SASRES (Tuukkanen *et al*., 2016) and SUPCOMB (prior to ATSAS version 3.1.0) (Kozin & Svergun, 2001).

Many of these programs are newly supported since the last publication about RAW. Users can now use RAW to align high resolution models with bead models via SUPCOMB/CIFSUP (depending on ATSAS version). They can do this either automatically when generating bead models as a final step in the process, or through the alignment window with any two saved models. Additionally, RAW now allows users to calculate theoretical scattering profiles from high resolution structures/models (either .pdb or .cif formats) using CRYSOL. If a user simply loads in a model the calculation is done automatically, or they can use the CRYSOL window to adjust the settings in CRYSOL, fit the model to measured experimental scattering data, and see a plot of the theoretical profiles and, if applicable, the normalized residuals to the experimental data. We have also added support for the new MW methods using Bayesian inference and machine learning, from the DATMW and DATCLASS programs, these are available in the MW window.

RAW is only tested against the latest version of ATSAS, so compatibility is only guaranteed for that version. However, attempts are made to preserve backwards compatibility, so previous versions will usually work, for example based on limited testing RAW 2.2.1 is compatible with ATSAS back to versions 3.0.X (which were 2-3 years old when this was written), as well as being fully compatible with 3.2.1 (the current latest release). Note that if backward compatibility breaking changes are made to the ATSAS programs between releases, RAW will not be compatible with the most recent ATSAS release until a new version of RAW is subsequently released.

### 4.7. Improvements to BIFT to add Monte Carlo error estimation and extrapolation of regularized curve to *I*(0)

We have updated the implementation of the BIFT method (Hansen, 2000) in RAW to provide Monte Carlo error estimation for the generated P(*r*) function and to extrapolate the regularized scattering profile to *I*(0). These updates coincided with moving the BIFT code from the depreciated scipy.weave compiler to the numba just-in-time (jit) compiler.

### 4.8. New documentation

For prior versions of RAW, we created the documentation manually in a Word document, saved it as a pdf, and uploaded it to the website with each new release of RAW. This was labour intensive, hard to update quickly, and awkward to actually use. As part of a complete overhaul of the documentation, we now create the documentation using the sphinx python package, and host it on Read the Docs. This allowed us to put the documentation in the program’s git repository, meaning it is now version controlled. With sphinx it is easy to build multiple different formats (including the html version on the web, and downloadable pdf, html, and epub versions), and updating it simply requires initiating a rebuild on Read the Docs, which grabs the latest version from the git. Read the Docs also provides multiple versions simultaneously. The latest version is the default, but every version of the documentation from the latest back to 1.3.1 is available (note that documentation versioning matches that of the RAW releases). The documentation is also distributed in HTML format with the RAW software, which is accessible on Windows and MacOS systems via the in-program Help menu.

We use the autodoc function in sphinx to automatically generate the API documentation for RAW, using the docstrings written as part of the API. In addition to tutorials in how to use the program itself, the website also has tutorials on best practices for basic SAXS operations like doing Guinier fits, calculation molecular weight, generating P(*r*) functions via IFTs, and making bead model reconstructions (https://bioxtas-raw.readthedocs.io/en/latest/saxs_tutorial.html).

In addition to the written documentation, most tutorials are now accompanied by YouTube videos (https://youtube.com/playlist?list=PLm39Taum4df4alFnacOOr1RWgylwiTWED), 23 in total with roughly 3.5 hours of runtime. These videos provide a useful alternative to the written tutorials for those who learn best by seeing the program in action. Finally, there is also a new (since the last report about RAW) mailing list where announcements about new versions are made and users can report bugs or ask questions (https://groups.google.com/g/bioxtas_raw).

## 5. New features

### 5.1. Comparison window

Since the previous RAW publication, we have added a dedicated window for easy comparison of multiple scattering profiles. The comparison window allows the user to compare profiles using the residual between profiles, the ratio of the profiles, or with a statistical test (currently the CORMAP (Franke *et al*., 2015) test implemented in RAW). For both residual and ratio comparisons the selected profiles are compared against a user selected reference profile. The user is shown a two-panel plot with all of the selected profiles plotted on the top and the appropriate metric plotted on the bottom. The user may then toggle profiles on or off, change which profile is the reference profile used for comparison, change the scaling of the plot axes, scale the profiles to the reference profile, and for the residuals plot display uncertainty normalized or un-normalized residuals. For the similarity test, each selected profile is compared against all the other selected profiles. The results are shown as a heatmap plot of the p value from the test, and in a list that gives the pair of compared profiles and the test result and p value. The results of the similarity test can be exported to a comma separated value (csv) file.

### 5.2. PDF reports

A major new feature in RAW is the ability to save a summary of your data processing for individual profiles, P(*r*) functions, and series data as a pdf report. A summary of DAMMIF/N and DENSS results produced by RAW can also be included. The report provides summary plots and tables of the key results, along with detailed tables of experimental parameters (when available) and all of the analysis results. The report is flexible, in that any or all of the data types can be saved in the same report, and more than one of each can be saved, in which case results are plotted on the same plots and added as additional columns to tables. Plots are saved as vector graphics, and so are suitable for inclusion in a publication (anecdotally, a number of these summary plots have been seen ‘in the wild’, usually in supporting information in lieu of the authors making their own versions). The window for generating the report makes it easy for the user to select which datasets loaded into RAW are included in the report by simply checking and unchecking boxes.

The summary plots saved in the report include: a plot of series intensity vs. frame number and *Rg* vs. frame number with selected buffer and sample ranges indicated, a log-lin plot, a Guinier plot with fit line and residual, a dimensionless Kratky plot for profiles and an *I*(0) normalized P(*r*) plot for P(*r*) functions. If the user did deconvolution on a series with EFA or REGALS then summary deconvolution plots are also saved. The summary table includes top line information such as *Rg* and *I*(0) values, MW values and *Dmax* values. Analysis specific tables provide more details, for example the Guinier analysis table provides the *q* range used for the Guinier fit, the *qmin*Rg* and *qmax*Rg* values, and the r^2^ of the fit, in addition to the *Rg* and *I*(0) values. A table of experimental parameters, such as date, experiment type, sample and buffer information, temperature, loaded volume and concentration, detector used, wavelength, etc is generated when these values are known. This typically requires that the reduction from images to 1D profiles be done in RAW, and that the beamline includes specific keywords in their metadata that RAW can recognize.

### 5.3. REGALS

Regularized Alternating Least Squares (REGALS) is a new method for deconvolving mixtures from SAS datasets (Meisburger *et al*., 2021). It can be applied to deconvolving overlapping peaks in SEC-SAXS, changing baseline and elution peaks in ion exchange chromatography SAXS, mixtures in time resolved SAXS data and equilibrium titration series data, and likely in other cases which we have not explored. We have created a GUI for REGALS in RAW that allows for fast, interactive deconvolution and provides initial SVD analysis of the entire dataset, evolving factor plots and SVD analysis of the background outside of the peak range (for chromatography data) to help the user determine how many components to use and where to start and end each range. Because the REGALS code is open source, REGALS is distributed with RAW and no additional installation is necessary for users (unlike the ATSAS programs) and changes based on the incorporation of REGALS into RAW have been pushed back to the original REGALS codebase.

Though the mathematics are not the same, from a practical point of view it can be thought of as an extension of the previously implemented evolving factor analysis (EFA) technique (Meisburger *et al*., 2016) to more complex conditions where components are not necessarily entering and exiting the dataset in a strict first-in/first-out approach like in SEC-SAXS. We still generally recommend EFA for standard SEC-SAXS data due to ease of use, but for more complex data as listed above, REGALS is preferred. REGALS can also handle deconvolution of SEC-SAXS data with a sloping baseline, something that EFA tends to fail at.

### 5.4. DENSS

Relatively recently, a new algorithm for determining actual (low resolution) electron density from solution scattering data (DENSS) was developed (Grant, 2018). This provides an alternative to the more traditional bead models. RAW provides users with a simple, user-friendly GUI interface for running DENSS, including creation of a number of individual density maps, averaging and refinement to create a final map, alignment of high-resolution structural models to the final density map, and evaluation of the results. The interface is very similar to the GUI for making bead models in RAW, making it easy for users to try either or both approaches for their data. Because the DENSS code is open source, DENSS is distributed with RAW and no additional installation is necessary for users (unlike the ATSAS programs) and changes based on the incorporation of DENSS into RAW have been pushed back to the original DENSS codebase.

## 6. The RAW API

A new capability for RAW is the ability to use it via an API mode without the associated GUI. In this case, the user installs RAW as a standard python package and then imports it into a standalone python script, interactive terminal session (e.g. iPython), Jupyter notebook, or another python program. The API provides access to all of the data reduction and analysis tools and algorithms available in the GUI, as well as direct access to the underlying experimental data objects that can be manipulated and saved. Profiles, P(*r*) functions, and series generated in the GUI can be loaded by the API, and vice versa for complete cross compatibility. The API is fully documented and several usage examples are provided as part of the RAW documentation (https://bioxtas-raw.readthedocs.io/en/latest/api.html).

Having a headless version of RAW has several significant use cases. The first is that it can now be used to process data in a much more automated fashion than the GUI allows, something that we have taken advantage of in the beamline processing pipeline at the BioCAT beamline (https://github.com/biocatiit/saxs-pipeline). The second is that it can be used to create custom processing scripts for data that cannot be easily handled in the GUI, for example in cases where the number of profiles make manual processing prohibitive or the dataset is not structured in a way that the GUI can handle. One example of this that fits both criteria is the time resolved SAXS data set used as part of (Martin, Harmon *et al*., 2021). The initial dataset involved ∼220,000 images (across all the conditions and repeats), which were reduced, averaged, and analysed initially by custom scripts using the API to yield the several hundred final profiles (over several different time series) that were then analysed by a combination of automated and manual approaches. The reduction and initial analysis scripts using the API are available as part of the supporting information of the referenced work (note: the scripts were written for an older version of the API, and may not run in the current version without modification).

## 7. Technical updates

We have made some significant technical changes to the RAW program since the previous publication. The most important of these was the update to Python 3 (Python Software Foundation, https://www.python.org/) compatibility and the replacement of the depreciated scipy.weave compiler with the numba just-in-time compiler for code that needed significant speed improvements. Both of these updates were critical for the ongoing maintenance and development of RAW and the ease of distribution of pre-packaged versions. We routinely update RAW to maintain combability with the latest version of the Python packages used by the program. As RAW also provides a GUI for various programs in the ATSAS package (requires separate installation) (Manalastas-Cantos *et al*., 2021) we also do routine maintenance to retain compatibility with new versions of ATSAS. Additionally, we moved RAW from a svn to a git for version control of the code and migrated the code from Sourceforge (still used to host release files) to GitHub.

One additional advantage of the creation of the RAW API is that it became possible to use pytest to create a large test suite with relatively comprehensive coverage for all non-GUI functionality in RAW (i.e. an analysis function can be automatically tested, but the GUI window that runs the analysis has to be manually tested). This is important to ensure functionality remains the same between versions and only intended changes occur. We test RAW on Windows, MacOS and Linux before each release.

As of this publication, the explicit package dependencies of RAW are: cython (https://cython.org/), dbus-python (Linux only) (https://dbus.freedesktop.org/doc/dbus-python/), fabio (Knudsen *et al*., 2013), future (https://python-future.org/), h5py (https://www.h5py.org/), hdf5plugin (http://www.silx.org/doc/hdf5plugin/latest/), matplotlib (Hunter, 2007), mmcif_pdbx (https://mmcif-pdbx.readthedocs.io/en/latest/), numba (Lam *et al*., 2015), numpy (Harris *et al*., 2020), pillow (https://pillow.readthedocs.io/en/stable/index.html), pyFAI (Ashiotis *et al*., 2015; Kieffer *et al*., 2019), reportlab (https://www.reportlab.com/), scipy (Virtanen *et al*., 2020), six (https://six.readthedocs.io/), svglib (https://github.com/deeplook/svglib) and wxpython (https://wxpython.org/). Additionally, we use sphinx (https://www.sphinx-doc.org/en/master/) and the sphinx_rtd_theme (https://github.com/readthedocs/sphinx_rtd_theme) to build the documentation, pytest (https://github.com/pytest-dev/pytest) for testing, and pyinstaller (https://pyinstaller.org/) to make the pre-built binaries. Note that most of these packages in turn depend on many other packages in the Python ecosystem.

## 8. Conclusions

RAW is a free open-source program that provides a wide range of basic and advanced analysis techniques for small angle scattering. Recent improvements include significantly expanded series processing capabilities, new automated methods for Guinier fitting and *Dmax* determination, ability to do electron density reconstructions, generation of PDF reports, a new API, and migration to Python 3. Designed to be an easy to learn program that can carry users from images through subtracted scattering profiles, model free analysis, and some basic model-based analyses, we believe the RAW package has a vital place in the SAS data analysis ecosystem, and that it will continue to prove useful for users for years to come.

## Acknowledgements

We thank the various people who contributed to code either in previous versions of RAW or in tools used by RAW, including: Soren Skou, Richard Gillilan, Thomas Grant, Steve Meisburger and Darren Xu. We would also like to thank Normand Cyr and Robert Miller who, along with Richard Gillilan, were beta testers for the python 3 conversion of RAW. We thank Nicholas Noinaj for his permission to use data collected by his group as anonymized examples in this paper. We thank Maxwell Watkins and Thomas Irving for providing feedback on the manuscript. This research used resources of the Advanced Photon Source, a U.S. Department of Energy (DOE) Office of Science User Facility operated for the DOE Office of Science by Argonne National Laboratory under Contract No. DE-AC02-06CH11357. BioCAT was supported by grant P30 GM138395 from the National Institute of General Medical Sciences of the National Institutes of Health. The content is solely the responsibility of the authors and does not necessarily reflect the official views of the National Institute of General Medical Sciences or the National Institutes of Health.

