## Supporting Information for "BioXTAS RAW 2: new developments for a free open-source program for small angle scattering data reduction and analysis"

Jesse B. Hopkins[a]

[a] The Biophysics Collaborative Access Team (BioCAT), Department of Physics, Illinois Institute of Technology, Chicago, IL 60616, USA

**Synopsis** BioXTAS RAW is a free, open-source program for reduction, analysis, and modelling of small-angle scattering data. This article describes the new and improved features in RAW version 2, including new tools for liquid chromatography coupled data processing, advanced reporting capabilities, and a new API.

**Synopsis** BioXTAS RAW is a free, open-source program for reduction, analysis and modelling of biological small angle scattering data. Here, the new developments in RAW version 2 are described. These include: improved data reduction using pyFAI; updated automated Guinier fitting and $D_{max}$ finding algorithms; automated series (e.g. SEC-SAXS) buffer and sample region finding algorithms; linear and integral baseline correction for series; deconvolution of series data using REGALS; creation of electron density reconstructions via DENSS; a comparison window showing residuals, ratios, and statistical comparisons between profiles; and generation of PDF reports with summary plots and tables for all analysis. In addition, there is now a RAW API, which can be used without the GUI, providing full access to all of the functionality found in the GUI. In addition to these new capabilities, RAW has undergone significant technical updates, such as adding Python 3 compatibility, and has entirely new documentation available both online and in the program.

#### S1. Automated series buffer and sample region selection

RAW's basic approach to finding a good buffer range is to scan a window of defined size along the measured profiles, and test each range (described below) to see if it is a valid buffer range. If no range is found, the window size is narrowed and the scan repeated until either a valid range is found or the minimum size is reached. RAW constrains the set of buffer ranges to test, both to avoid false positives and to improve the speed of the algorithm. Initially, it uses a peak finding algorithm (the *find_peaks* function in scipy) on a smoothed version of the intensity vs. frame data, which provides the position of all peaks in the dataset. If no peaks are found in the data the algorithm starts the search from the first frame (earliest point in the elution) and proceeds from there. Assuming peaks are found, this defines the starting search range and window size. The initial window size is twice the width of the largest peak at 40% of its maximum intensity above the baseline. The initial search range starts to left (early frame/time) of the first peak, and proceeds towards the start of the dataset (earliest frame/time) in a series of steps whose size depends on the size of the window. This prioritizes buffer measurements closer to the elution peak. For example, if RAW found a single peak at frame 100 with width 10 at 40% maximum intensity, the initial search window would be 20, and the step size would be 4. The ranges tested would be: 75-95, 71-91, 67-87, and so on until either a valid buffer range is found or the start of the dataset is reached.

If a valid buffer range is not found for the initial window size, RAW narrows the window size and redoes the search. This is repeated until a valid range is found or a defined minimum window size is reached. If a valid range has not been found once the minimum window size is reached, the algorithm then searches for a buffer range in the data collected after the last elution peak, using the same range of window sizes and again starting with ranges closer to the peak and then testing those further away. If a buffer range still is not found, RAW then searches ranges between the peaks.

The test for a valid buffer range has two inputs. The first is the total intensity (or mean intensity or intensity in a given $q$-range or at a particular $q$ value, depending on user choice) vs. frames (or time) data, sometimes called the scattergram, and the second is the scattering profiles at each measured point in the elution. RAW evaluates a buffer range in three ways. In the first part of the test it calculates the Spearman rank-order correlation coefficient and associated p value for the intensity vs. frame and smoothed intensity vs. frame data in the specified range. The p value, though not technically valid for small ranges, is an indicator of whether there are correlations in the intensity. Buffer scattering should have the same intensity at all measured points, so correlations are indicative of something eluting in the data (or an issue with the baseline), and RAW marks ranges with possible correlations as not valid.

In the second part of the test RAW checks the similarity of the scattering profiles in the selected range compared to the scattering profile with the median total intensity in the range, using the CORMAP test. The algorithm tests three different $q$ ranges: the full $q$ range, the low q range (first 100 points) and the high $q$ range (last 100 points). A p value calculated for each range determines if all profiles are similar across each tested q range. RAW uses three different *q* ranges to test for different artifacts in the data. Changes in the low-*q* may be indicative of capillary fouling or other unwanted damage effects on the data. Changes in the high-*q* may come from beam or temperature drift (though these can also show up at low-*q*), while changes in the full profile acts as a catch-all metric. Because CORMAP relies on the number of outlier points vs. the total number of points to generate a p value, doing the test on a smaller number of points makes it more sensitive to a few outliers, such as a small number of points changing at the lowest *q* values that might indicate capillary fouling. RAW marks ranges where any of the buffer profiles are different from the median profile in any of these three *q* ranges as not valid.

In the third part of the test, RAW performs a singular value decomposition (SVD) on the entire selected set of profiles and find the number of significant singular values. Buffer ranges should only have one significant singular value, so if there is more than one significant singular value in the tested range than it is not a valid buffer range. In our experience this is typically the least sensitive test, but as RAW has the capability already available it is included for completeness.

In order to optimize the speed of the automated buffer finding, the RAW runs the parts of the test in the order listed above, from fastest to slowest, and if any part fails on the selected range RAW does not run the subsequent parts.

RAW uses the same general approach for the automated sample range determination as it does for the automated buffer range finding, a window is scanned along the data and it tests whether each range is a valid sample range (described below). RAW constrains the sample ranges to test, both to avoid false positives and to improve the speed of the algorithm. The midpoint of the largest peak is selected as the starting point, and an initial search window size is set equal to the width of the peak at 40% of the maximum intensity above baseline. A search range is defined as twice that width. The search window is then shifted alternatively to earlier in the elution and then later in the elution, with a step size for the shift based on the window size, with each alternation getting further from the midpoint. For example, if the midpoint was 100, the window size 10, and the shift step size 2, the windows tested would be: 95-105, 93-103, 97-107, 91-101 and so on until a valid range was found. If no valid range is found for the initial window size, the window size is reduced and the range retested until either a valid range is found or the minimum range size is reached and the algorithm fails to find a valid sample range. A valid sample range will not be found if no peaks are found in the dataset.

The test for a valid sample range has three inputs: the scattering profiles and the $R_g$ and MW calculated for each profile in the selected range. There are five parts to the test. The first part is simply whether all profiles in the selected range have calculated $R_g$ and MW values. If the values could not be calculated for all profiles in the range, then the range is not a valid sample range. The second part calculates the Spearman correlation coefficient and p value for the $R_g$ and MW values in the range. If the sample is uniform across the selected range there should be no correlation, so if the p value from this test indicates a correlation, RAW marks the range as not valid.

The third part tests for similarity between the subtracted scattering profiles in the selected range and the subtracted scattering profile with the maximum total intensity in that range, using the CORMAP test. Because we expect that the intensity of the profiles will change across the peak due to the changing concentration in elution, RAW scales all profiles to the profile with the maximum intensity before the similarity test is done. As with the buffer similarity test, the full $q$ range, the low-$q$ range, and the high-$q$ range are all tested, and if there are any profiles that are not similar to the highest intensity profile in any of the three $q$ ranges then RAW marks the sample range as not valid.

For the fourth part, RAW performs a SVD on the entire selected set of profiles and find the number of significant singular values. As with the buffer range SVD test, subtracted sample ranges should only have one significant singular value, so if there is more than one significant singular value in the selected range than it is not a valid sample range. Again, this tends to be the least sensitive test, but we include it because it was already available in RAW.

The fifth and final part of the test is to check whether including all the profiles in the selected range improves the signal to noise of the final averaged subtracted scattering profile. Here, RAW sorts the profiles in the range by their overall intensity. An average subtracted profile is created by starting with just the most intense profile, and then subsequently averaging that with the next most intense profile, and so on. Every time a new profile is included in the average, RAW calculates the mean of the intensity/uncertainty in the average profile across all $q$ points, yielding the overall signal to noise ratio. If that signal to noise ratio decreases when a profile is included, then that profile should not be included in the final dataset for optimal signal to noise, and so the selected range is not valid.

As with the automated buffer range selection, in order to optimize the speed of the algorithm RAW runs the tests in the order listed above, fastest to slowest, and if any test fails on the range subsequent tests are not run.

#### S2. Further information on automated $R_g$ and $D_{max}$ determination

##### S2.1. Automated $D_{max}$ determination

The auto $D_{max}$ function can run in several ways. If the ATSAS package is not available, it simply returns the $D_{max}$ value found by BIFT. However, if the ATSAS package is available then $D_{max}$ can be fine-tuned to get a more accurate value. The basic idea is simple. First, RAW runs other automated methods – BIFT, DATGNOM (Petoukhov *et al.*, 2007) and DATCLASS – to determine a good starting point for the search. Based on the results from the SASBDB when the algorithm was written, similar to the evaluation below, if the DATCLASS $D_{max}$ is available RAW uses it as the starting point. If not, RAW uses the average of the BIFT and DATGNOM $D_{max}$. If only the BIFT or DATGNOM $D_{max}$ is returned (but ATSAS is available), RAW applies a compensation factor for the observed overestimation and underestimation (below), giving a starting $D_{max}$ of 0.79*(BIFT $D_{max}$) or 1.2*(DATGNOM $D_{max}$). After determining a starting value, RAW calculates the P($r$) function using GNOM with force to zero at the maximum dimension turned off. RAW checks this initial unconstrained P($r$) function in two ways, first for overestimated $D_{max}$ values then for underestimated $D_{max}$ values, and adjusts the maximum value, as described below, until it finds a good value.

In a P($r$) function with an overestimated $D_{max}$, we expect either a long tail oscillating about zero (for homogenous, monodisperse, non-interacting samples) or negative values near the maximum dimension (for data with repulsive interactions) (Jacques & Trewhella, 2010). RAW uses two mechanisms to check for this and adjust the $D_{max}$ value. First, if any of the last 20 values of the P($r$) function are less than zero RAW marks the value as overestimated. In that case, RAW decreases $D_{max}$ in 1 Å increments and recalculates the unconstrained P($r$) function using GNOM until this criterion is no longer satisfied. Then RAW checks the value of the P($r$) function at $D_{max}$, and if it is less a defined threshold fraction of the maximum value of the P($r$) function (by default 1%, but can be adjusted in the API) it decreases $D_{max}$ by 1 Å and recalculates the P($r$) function until that threshold is met. These two checks and adjustments prevent significant overestimates of $D_{max}$.

In a P($r$) function with an underestimated $D_{max}$, we expect that the value at the end of the P($r$) function is significantly greater than zero (Jacques & Trewhella, 2010). If the value of the P($r$) function at $D_{max}$ is greater than a defined threshold fraction of the maximum value of the P(r) function (by default 1%, but can be adjusted in the API) RAW increases $D_{max}$ by 1 Å and recalculates the P($r$) function until that threshold is met. This prevents significant underestimates of $D_{max}$.

RAW applies one additional constraint to the adjustments, constraining the change in $D_{max}$ to be no more than 50%, either an increase or decrease, of the initial value, to prevent the algorithm from running away. This is particularly useful in the cases of highly aggregated data where there may be no appropriate maximum dimension and the algorithm could otherwise increase $D_{max}$ essentially indefinitely.

When taken together, these two simple adjustments for overestimated and underestimated $D_{max}$ values provide a more robust estimate of the maximum dimension than any of the other tools mentioned above, though we rely on those tools to find an appropriate starting point and so our approach should be considered complementary to the previously developed methods.

#### S2.2. Comparison of automated $R_g$ determination

Using the approach described in the main text, we compared automated methods for determining $R_g$ against experimenter reported values in the SASBDB. Here we report on the results when all models are included, even ones where the $R_g$ reported may be directly from an automated method. In this case, 3110 of the initial 3138 datasets had experimenter reported $R_g$ values. From these, the average and standard deviation of the ratio of (experimenter determined $R_g$)/(automatically determined $R_g$) was 1.03 ± 0.49 for the RAW auto Guinier method and 1.02 ± 0.48 for the ATSAS AUTORG method. Plots of the automated $R_g$ vs. experimenter determined $R_g$ are shown in Figure S2. RAW failed to return results for 8 (0.26%) datasets and AUTORG failed to return results for 16 (0.51%) datasets. These results again show that both algorithms are robust, in that they fail on less than 1% of all datasets, and that both algorithms are on average quite accurate. As before, the old RAW algorithm, run on the same set of data, was similarly accurate ($R_g$ ratio: 1.02 ± 0.44), but failed on a significant number of datasets (362, 11.5%).

The results can be further broken down by molecule type. Table S1 gives the results for the three different available categories: Protein, RNA, and DNA. All available datasets were analysed. Table S2 shows the same thing, but excludes results where the experimenter provided *q*-range matched that determined by one of the automated methods.

#### S2.3. Comparison of automated $D_{max}$ determination

Using the approach described in the main text, we compared automated methods for determining $D_{max}$ against experimenter reported values in the SASBDB. Here we report on the results when all models are included, even ones where the $D_{max}$ reported may be directly from an automated method. In this case, 2875 of the initial 3138 datasets had experimenter reported $D_{max}$ values. From these, the average and standard deviation of the ratio of (experimenter determined $D_{max}$)/(automatically determined $D_{max}$) was determined for four methods: RAW auto $D_{max}$, the ATSAS DATGNOM and DATCLASS methods, and the BIFT method implemented in RAW. The results are: 0.90 ± 0.48 for the RAW auto $D_{max}$ method, 1.25 ± 0.92 for DATGNOM, 1.05 ± 0.52 DATCLASS, and 0.83 ± 0.65 for BIFT. Plots of the automated $D_{max}$ vs. experimenter determined $D_{max}$ are shown in Figure S3. RAW, DATGNOM, and BIFT all returned results for every dataset, whereas DATCLASS failed to return results for 536 (18.6%) datasets. These results show that none of the algorithms are both robust (returning results for most/all datasets) and accurate. DATCLASS has the best average accuracy (and second lowest standard deviation), but fails on a large fraction of the datasets. RAW's auto $D_{max}$ approach is the next best, which is not surprising since it incorporates information from the other three, but tends to overestimate $D_{max}$ relative to the experimenter provided values. BIFT is somewhat less accurate than RAW and also overestimates $D_{max}$, while DATGNOM is the least accurate and tends to significantly underestimate $D_{max}$ while also having the largest deviation.

The results can be further broken down by molecule type. Table S3 shows the results for the three different available categories: Protein, RNA, and DNA. All available datasets were analysed. Table S4 shows the same thing, but excludes results where the experimenter determined value matched that determined by one of the automated methods.

We also want to note that a question can and should be raised about the quality of the experimenter provided values. Finding an appropriate $D_{max}$ value is significantly harder than getting a good Guinier fit, and the general rule of thumb in the community is that for good quality data, experienced workers can find $D_{max}$ values that are different by up to 10% and still have reasonable results. It is even harder to find appropriate values for non-ideal datasets, either those that are not monodisperse or those exhibiting structure factor effects in the scattering profile. This reality is reflected in the less accurate automated methods when compared with the automated Guinier fitting. Our personal, anecdotal, experience in working with RAW to analyse data almost every day is that the automated method, for good quality data, tends to be quite good, and so we wonder if there may be some amount of routine underestimation in reported $D_{max}$ values. Because pursuing this line of questioning more thoroughly would require manual analysis of thousands of datasets it is beyond the scope of this paper.

#### S3. General data collection method

Some data presented in this paper was collected by users at the BioCAT beamline, where we are based, and has been anonymized and used with permission for the purposes of showing various characteristic behaviours of automated algorithms. The goal of the paper is not to analyse this data, and so only a general method for the collection is presented, specific details (such as exact exposure times, sample to detector distance, etc) may be slightly different between the datasets. For the same reason this data is not made available for further analysis.

SAXS was performed at BioCAT (beamline 18ID at the Advanced Photon Source, Chicago) with in-line size exclusion chromatography (SEC-SAXS) to separate sample from aggregates and other contaminants thus ensuring optimal sample quality. Sample was loaded onto a Superdex 200 Increase 10/300 GL column (Cytiva), which was run at 0.6 ml/min by an AKTA Pure FPLC (Cytiva) and the eluate after it passed through the UV monitor was flown through the SAXS flow cell. The flow cell consists of a 1.0 mm ID quartz capillary with ~20 μm walls. A coflowing buffer sheath is used to separate sample from the capillary walls, helping prevent radiation damage (Kirby *et al.*, 2016). Scattering intensity was recorded using an Eiger2 XE 9M (Dectris) detector which was placed 3.6 m from the sample giving us access to a *q*-range of 0.003 $Å^{-1}$ to 0.42 $Å^{-1}$. 0.5 s exposures were acquired every 1 s during elution and data was reduced using BioXTAS RAW (Hopkins *et al.*, 2017).

**Table S1** The success of the automated $R_g$ determination methods by molecule type as listed in the SASBDB. The average across all molecule types ("All") is included for reference.

| Type | Datasets with $R_g$ values | RAW $R_g$ ratio | RAW failures | ATSAS $R_g$ ratio | ATSAS failures | Old RAW $R_g$ Ratio | Old RAW failures |
|---|---|---|---|---|---|---|---|
| All | 3110 | 1.03±0.49 | 8 (0.26%) | 1.02±0.48 | 16 (0.51%) | 1.02±0.44 | 362 (11.5%) |
| Protein | 2770 | 1.03±0.48 | 8 (0.29%) | 1.01±0.46 | 13 (0.47%) | 1.02±0.45 | 300 (10.8%) |
| RNA | 216 | 1.08±0.64 | 0 (0%) | 1.01±0.08 | 1 (0.46%) | 1.01±0.07 | 45 (20.8%) |
| DNA | 124 | 1.07±0.52 | 0 (0%) | 1.14±1.06 | 2 (1.6%) | 1.06±0.55 | 17 (13.7%) |

**Table S2** The success of the automated $R_g$ determination methods by molecule type as listed in the SASBDB, excluding datasets where the experimenter input $q$-range matched one of the $q$-ranges returned by the RAW or ATSAS automated method. The average across all molecule types ("All") is included for reference.

| Type | Datasets with $R_g$ values | RAW $R_g$ ratio | RAW failures | ATSAS $R_g$ ratio | ATSAS failures | Old RAW $R_g$ Ratio | Old RAW failures |
|---|---|---|---|---|---|---|---|
| All | 1827 | 1.04±0.56 | 5 (0.27%) | 1.03±0.63 | 11 (0.60%) | 1.02±0.51 | 190 (10.4%) |
| Protein | 1578 | 1.04±0.58 | 5 (0.32%) | 1.02±0.60 | 8 (0.51%) | 1.02±0.52 | 146 (9.3%) |
| RNA | 162 | 1.05±0.38 | 0 (0%) | 1.01±0.10 | 1 (0.62%) | 1.02±0.08 | 33 (20.4%) |
| DNA | 87 | 1.09±0.61 | 0 (0%) | 1.20±1.26 | 2 (2.3%) | 1.08±0.65 | 11 (12.6%) |

**Table S3** The success of the automated $D_{max}$ determination methods by molecule type as listed in the SASBDB. The average across all molecule types ("All") is included for reference.

| Type | All | Protein | RNA | DNA |
|---|---|---|---|---|
| Datasets with $D_{max}$ values | 2875 | 2553 | 203 | 119 |
| RAW $D_{max}$ ratio | 0.90±0.48 | 0.89±0.46 | 1.02±0.66 | 0.90±0.51 |
| RAW failures | 0 (0%) | 0 (0%) | 0 (0%) | 0 (0%) |

| | | | | |
|---|---|---|---|---|
| DATGNOM $D_{max}$ ratio | 1.25±0.92 | 1.25±0.97 | 1.25±0.35 | 1.31±0.55 |
| DATGNOM failures | 0 (0%) | 0 (0%) | 0 (0%) | 0 (0%) |
| DATCLASS $D_{max}$ ratio | 1.05±0.52 | 1.05±0.53 | 1.04±0.35 | 1.05±0.24 |
| DATCLASS failures | 536 (18.6%) | 409 (16.0%) | 84 (41.4%) | 43 (36.1%) |
| BIFT $D_{max}$ ratio | 0.84±0.65 | 0.84±0.67 | 0.77±0.36 | 0.90±0.60 |
| BIFT failures | 0 (0%) | 0 (0%) | 0 (0%) | 0 (0%) |

**Table S4** The success of the automated $D_{max}$ determination methods by molecule type as listed in the SASBDB, excluding datasets where the experimenter input $q$-range matched one of the $q$-ranges returned by the RAW or ATSAS automated method. The average across all molecule types ("All") is included for reference.

| Type | All | Protein | RNA | DNA |
|---|---|---|---|---|
| Datasets with $D_{max}$ values | 2502 | 2217 | 185 | 100 |
| RAW $D_{max}$ ratio | 0.89±0.51 | 0.88±0.49 | 1.02±0.69 | 0.90±0.55 |
| RAW failures | 0 (0%) | 0 (0%) | 0 (0%) | 0 (0%) |
| DATGNOM $D_{max}$ ratio | 1.27±0.98 | 1.27±1.03 | 1.26±0.36 | 1.36±0.59 |
| DATGNOM failures | 0 (0%) | 0 (0%) | 0 (0%) | 0 (0%) |
| DATCLASS $D_{max}$ ratio | 1.06±0.56 | 1.06±0.57 | 1.03±0.36 | 1.06±0.26 |
| DATCLASS failures | 501 (20.0%) | 382 (17.2%) | 82 (44.3%) | 37 (37.0%) |
| BIFT $D_{max}$ ratio | 0.84±0.69 | 0.84±0.71 | 0.78±0.35 | 0.90±0.64 |
| BIFT failures | 0 (0%) | 0 (0%) | 0 (0%) | 0 (0%) |

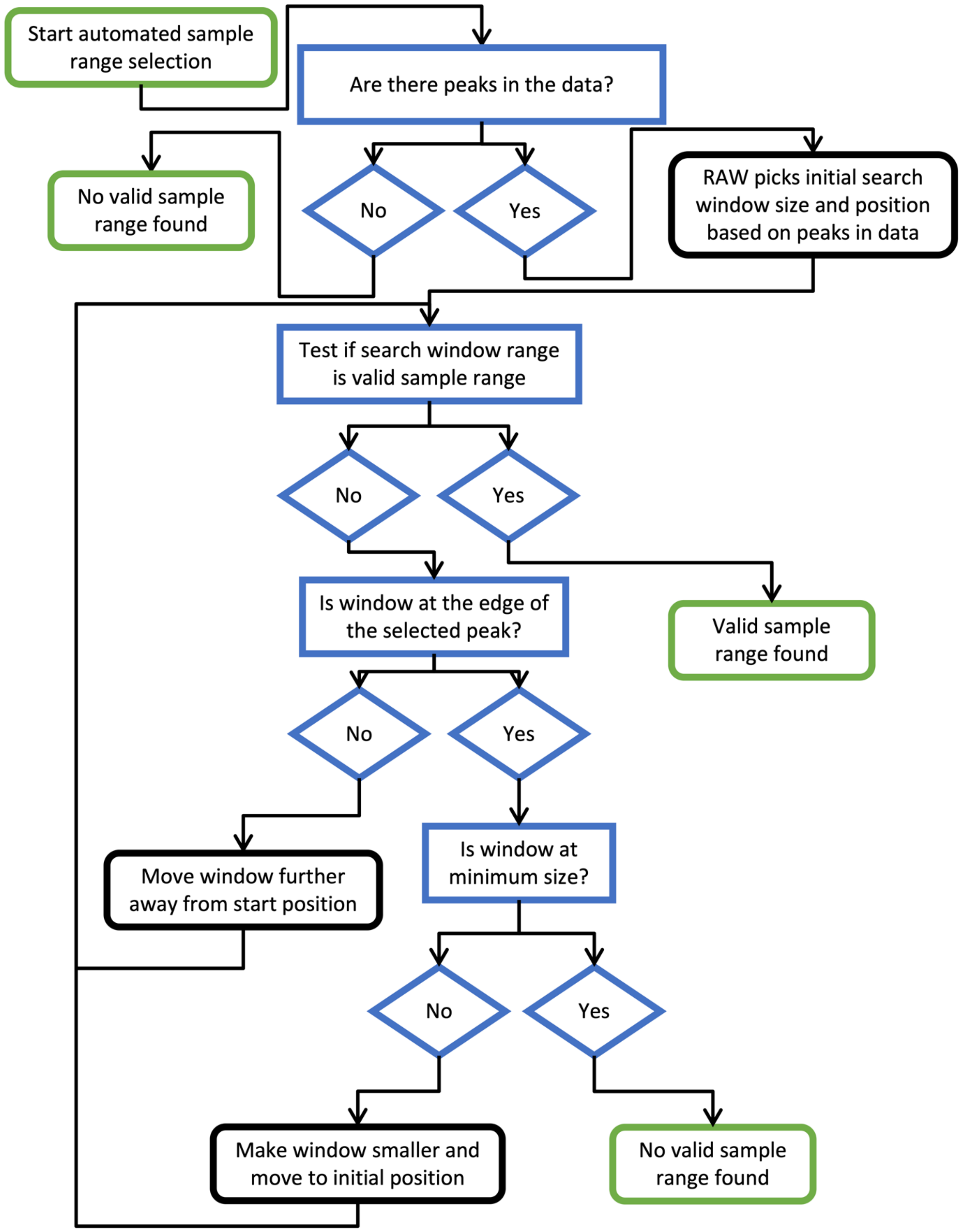


**Figure S1** A flow chart for the automated sample range finding algorithm used by RAW. Green edged boxes (rounded corners) are start and end points, blue edged boxes (square corners) and diamonds are decision points or tests in the algorithm, and black edged boxes (rounded corners) are actions by the algorithm.

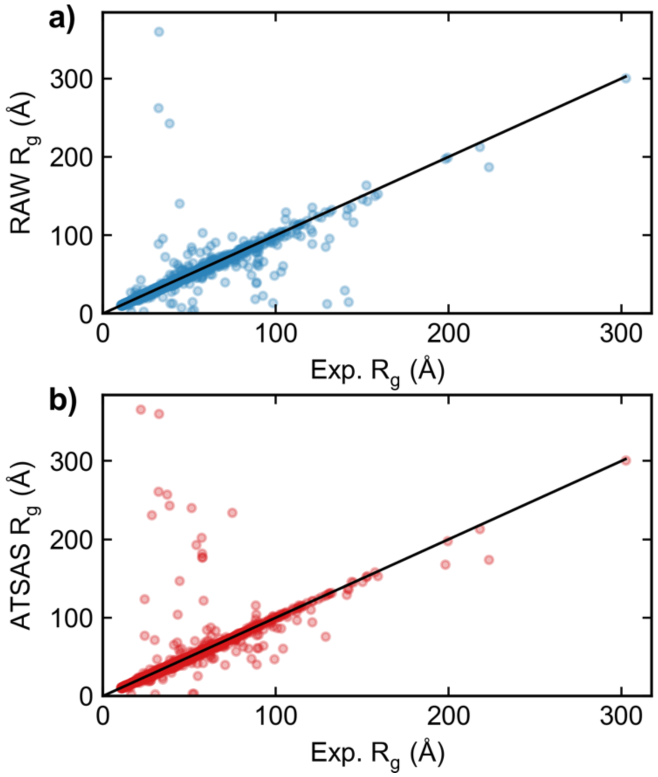


**Figure S2** Plots of the automatically calculated $R_g$ by a) the RAW automatic Guinier function, and b) the ATSAS AUTORG function on the y-axis vs. the experimenter determined $R_g$ from a SASBDB entry on the x axis. Results are shown for all SASBDB entries with $R_g$ values that were classified as either Protein, DNA, or RNA. Perfect agreement between the automated method and the experimental method would be equal $R_g$ values, shown by the black line in each figure.

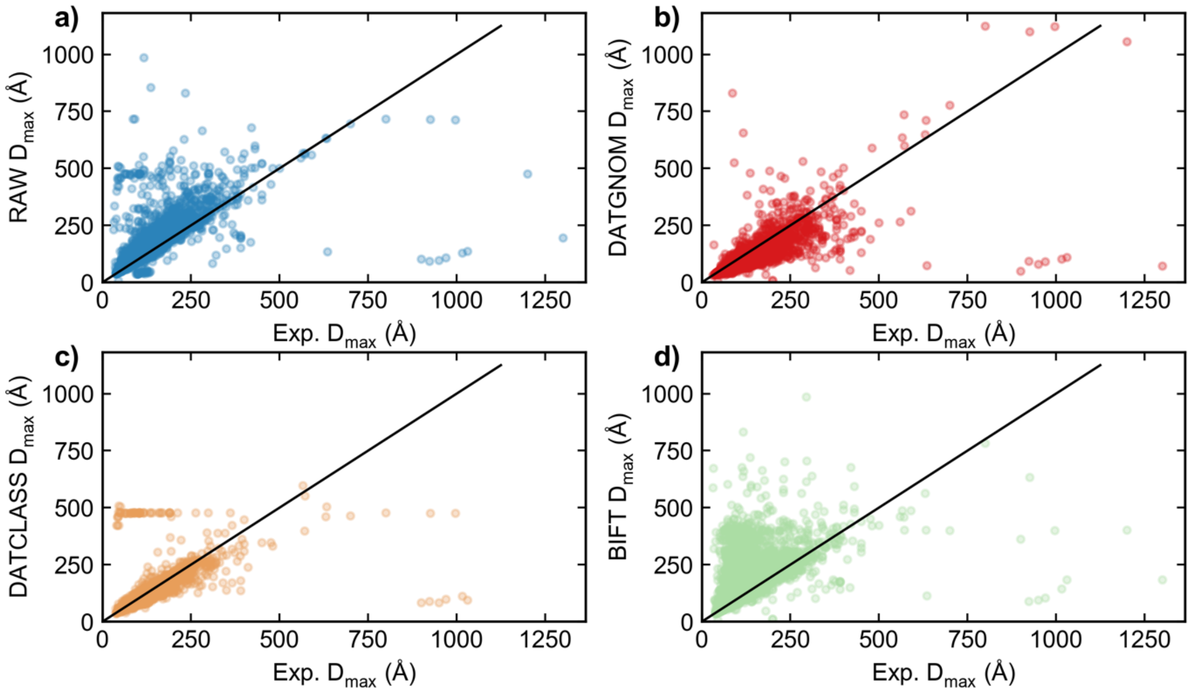


**Figure S3** Plots of automatically calculated $D_{max}$ by a) the RAW auto $D_{max}$ function, b) the ATSAS DATGNOM function, c) the ATSAS DATCLASS function, and d) BIFT (as implemented in RAW) on the y-axis vs. the experimenter determined $D_{max}$ from a SASBDB entry on the x axis. Results are shown for all SASBDB entries with $D_{max}$ values that were classified as either Protein, DNA, or RNA. Perfect agreement between the automated method and the experimental method would be equal $D_{max}$ values, shown by the black line in each figure.

### References

References are included in the reference list for the main text.
